# Semantic Dimensions Support the Cortical Representation of Object Memorability

**DOI:** 10.1101/2024.10.03.616545

**Authors:** Matthew Slayton, Cortney M Howard, Shenyang Huang, Mariam Hovhannisyan, Roberto Cabeza, Simon W Davis

## Abstract

Recent work in vision sciences contends that objects carry an intrinsic property, memorability, that describes the likelihood that an object can be successfully encoded and later retrieved from memory. It has been shown that object memorability is supported by semantic information, but the neural correlates involved are largely unexplored. The present study explores these premises and asks whether neural correlates of object memorability can be accounted for by semantic dimensions. To investigate these questions, we combine three datasets: (1) feature norms for a database of ∼1000 natural object images, (2) normative conceptual and perceptual memory data for those objects, and (3) neuroimaging data from an fMRI study collected using a subset (n=360) of those objects. We found that object-wise memorability elicits consistent brain activation across participants in key mnemonic regions (e.g., hippocampus and rhinal cortex), and that a substantial portion of the variance in this brain activity is mediated by the semantic factors describing these images. Regions with the strongest mediation effects are associated with sensory, motor, and visual processes, suggesting that the relationship between memorability and semantics may align with a sensory-functional account of the representation of concepts in the brain.

## INTRODUCTION

The growth of object-wise analysis in both psychology and computer vision science has led to a focus on prediction of object salience and memorability based on reliable stimulus properties. This focus on object-level properties has led many researchers to notice that some object images are better remembered than others when considered across participants (Isola et al., 2011, 2014), across a range of stimulus categories (Bainbridge et al., 2013), and across a range of testing conditions (Wakeland-Hart et al., 2022). While some perspectives on this phenomenon hold that object memorability constitutes an intrinsic attribute of visual stimuli (Bainbridge, 2022), there is growing evidence that supports an alternative yet complementary view that this phenomenon may be largely explained by visual and semantic features of the stimuli (Deng et al., 2024; Hovhannisyan et al., 2021; Kramer et al., 2023). Adjudicating between these two models of memorability can be achieved through neuroimaging which can answer explicit questions about whether memorability is associated with consistent cortical activity, and more importantly, whether this activity can be accounted for by properties of the stimuli. Therefore, the cognitive neuroscience of object memory is approaching a crossroads, especially concerning the role of semantic information in promoting episodic memory for visual images. The current study seeks to close the loop in this conversation, by using the brain as a model for investigating the utility of using semantic features to explore object memorability.

Several groups have identified factors that influence image memorability, including attentional state (Wakeland-Hart et al., 2022), image meaningfulness (Shoval et al., 2023), as well as the semantic and visual features of the images themselves (Bainbridge, 2022; Deng et al., 2024; Dubey et al., 2015; Hovhannisyan et al., 2021). The neural basis of the encoding of such features spans a rigorously explored hierarchy from early visual cortex, through the ventral stream towards more abstract processing in anterior temporal and frontal cortices (Peelen & Caramazza, 2012; Tyler et al., 2013). It is known that in visual processing, conceptual features are encoded in multiple regions along the ventral stream that serve numerous functional roles, such as category differentiation in fusiform gyrus (Tyler et al., 2013), feature differentiation in perirhinal cortex (Clarke & Tyler, 2014), and semantic representation in anterior temporal lobe (Chen et al., 2016). Several of these regions support the encoding and retrieval of visual stimuli, including early visual cortex, lateral occipital cortex, and superior temporal sulcus (Lahner et al., 2024), as well as perirhinal cortex, parahippocampal cortex, and fusiform gyrus (Bainbridge et al., 2017; Bainbridge & Rissman, 2018). The current analysis proceeds on the premise that neural representations of reliable conceptual features represented in the ventral stream drive the ability of humans to remember what they see.

While there appears to be consensus regarding the role of conceptual feature encoding in the brain and its impact on image memorability, there are important differences between the methodologies currently used to model image features. Some groups use normed image features (Davis et al., 2021; Devereux et al., 2013; Hovhannisyan et al., 2021) while others use image similarity judgments to derive semantic dimensions (Bainbridge, 2022; Contier et al., 2023; Hebart et al., 2020; Kramer et al., 2022, 2023). Kramer et al. set such a high-dimensional model against other potential explanations of image memorability, such as object typicality for a given dimension. They found that features are stronger predictors of memorability than object typicality, and that semantic features have a larger role than do perceptual features, explaining 88% of the variance in memorability (Kramer et al., 2023). Similarly, Hovhannisyan et al. found that complex semantic features (e.g., the distinctiveness of features for a given image) are generally much better predictors of image memorability than either simple semantics (e.g., image frequency) or the visual properties of the image (Hovhannisyan et al., 2021). These results inform our focus on semantic features in mediating the relationship between memorability and brain activity.

In the current study, we endeavored to build on this work by investigating the role that semantic information has in explaining the relationship between image memorability and image-related brain activity. We used a previously published neuroimaging dataset (Davis et al., 2021) using feature norms and memorability norms (Hovhannisyan et al., 2021) in order to build a model of image memorability in the brain. We further compared the relative roles of human-derived image memorability ratings and semantic dimensions based on stimulus features norms in predicting trial-wise brain activity. We expected that object-wise memorability would elicit consistent brain activation in key mnemonic regions (e.g., hippocampus and parahippocampal cortex), and that a substantial portion of the variance in this brain activity could be explained by the semantic factors underlying these images. The goal was to provide a concrete neural correlate of the image properties engendering the memorability effect.

## METHODS

### Methods Overview

The present study examined extant datasets from three related experiments which have been examined in previous behavioral analyses (Hovhannisyan et al., 2021) or neural basis of object memory (Davis et al., 2021; Huang et al., 2024). The first experiment collected a feature norming database based on online descriptions of a set of object images, the second conducted an online experiment testing conceptual memory for the same objects, and a third neuroimaging experiment used objects from this database to investigate the role of cortical representations in mediating conceptual memory. While the full description for these data can be found in the original publications (Davis et al., 2021; Hovhannisyan et al., 2021), here we describe details of study design, acquisition and analysis sufficient to understand the current study.

### Stimuli

The total set of stimuli include 995 object images drawn from a variety of object categories, including mammals, birds, fruits, reptiles, sea animals, vegetables, tools, clothing items, foods, musical instruments, vehicles, furniture items, buildings, and other objects found within specific cultural or functional contexts. The 12 object categories were balanced for frequency based on the Corpus of Contemporary American English (Davies, 2008). The full database of objects and features can be accessed at: https://mariamh.shinyapps.io/dinolabobjects/

### Experiment 1: Feature Norming

#### Data collection

566 participants were recruited via Amazon Mechanical Turk (all with greater than or equal to 95% approval) for the initial feature norming experiment. Based on self-identification, this population comprised 347 women and 219 men, mean age = 34.6 years (range 19–75), all self-reported native speakers of American English. Participants completed between one and five sessions which lasted one hour and included 40 presented images. Participants were paid $3.00 per hour for their participation in the property norming study. Informed consent was obtained from all participants under a protocol approved by the Duke Medical School Institutional Review Board (IRB). All procedures and analyses were performed in accordance with IRB guidelines and regulations for experimental testing.

Participants were shown an object (e.g., a porcupine) and were given a space to add five unique features, similar to previous feature-norming paradigms (Devereux et al., 2014; McRae et al., 2005). Participants were given a sentence (e.g., “Porcupine is/has ”) and would fill in a feature, such as “an animal” or “quills.” Stimuli were randomized across participants. No two images from the same category appeared consecutively, and each image was presented to at least 20 participants.

Features were then cleaned following the manual procedures used by McRae et al. (McRae et al., 2005) and Devereux et al. (Devereux et al., 2014), including: (1) removal of adverbs, (2) feature splitting, for example a feature such as “has a round face” was rewritten as “has a round face” and “has a face,” (3) combining close synonyms, for example replacing “groups, packs, and flocks” with “groups”, (4) correction of spelling mistakes, (5) morphological mapping, for example combining “is used in cooking” and “is used by cooks” into “is used in cooking,” (6) removal of plural forms, and (7) removal of features not present in at least two objects. Afterward, a feature × concept production frequency matrix was created to describe the normalized frequency of a given feature for a given concept. Each feature was further assigned a feature category label first used by McRae et al., (McRae et al., 2005), for example encyclopedic (e.g. “lives in the US,” *n* = 1930 features), visual (e.g., “is red,” *n* = 1886 features), and functional (e.g. “used for cutting,” *n* = 1101 features). These feature categories help to classify features, and there is high interrater reliability (ICC > 0.8).

Both human-derived and text corpus-derived methods are used for modeling and experimental investigations in human cognitive neuroscience, and consensus on which method is best for the study of semantics and human memory is still evolving. Feature norm approaches have a nearly twenty-year history in the field and have been used by several groups (Borovsky et al., 2023; Devereux et al., 2014; McRae et al., 2005; Van Overschelde et al., 2004; Vinson & Vigliocco, 2008). We chose a method with a history of use in cognitive neuroscience in order to facilitate comparison with previous results and maximize reliability.

### Experiment 2: Online Memory Testing

A separate group of participants (Amazon Mechanical Turk, >95% approval rating, all self-reported speaker of native English) participated in the conceptual (*n* = 200) recognition memory task. After excluding responses from 7 users due to computer error, the conceptual memory task included 193 users (108 women and 85 men, 19–87 years of age, mean = 39.7 years). This study used the same 995 objects used in the feature norming study. The AMT workers completed the encoding task on Day 1, determining if an object was living or non-living. Each image was presented for 2 s with a 1s inter-trial interval. On Day 2 (±4 h; mean lag between Encoding and Retrieval = 29.34 h) participants completed a two-alternative forced-choice oldness decision, whether an object was seen on Day 1 (i.e., “old”) or not (i.e., “new”). 168 old and 168 new stimuli were presented, balanced across object categories. Workers were paid $0.50 for completing the Day 1 encoding session (mean time to finish 16.07 min), and $4.50 for completing the Day 2 retrieval session (mean time to finish the tasks was 23.5 min). After data collection, mean hit rates and false alarm rates for each object were calculated based on the percentage of correct responses across subjects to old or new trials, giving object-wise average memorability ratings. Corrected Recognition rates (Hit Rate – False Alarm Rate) were calculated for each object.

### Experiment 3: Neuroimaging

Details for the neuroimaging experiment have been published previously (Davis 2021, Huang 2024, Howard 2024). In brief, 26 subjects were recruited (native English speakers, 14 women, age mean ± SWD 20.4 ± 2.4 years; range 18–26 years) in accordance with a protocol approved by the Duke University Health System Institutional During the study, each object was presented alone on white background in the center of the screen. Subjects performed memory tasks across two separate days: Day 1 for encoding and Day 2 for retrieval. On Day 1, they viewed object images and were asked to covertly name the image. They also pressed a button to indicate that the image matched a single letter probe shown immediately before (e.g. a letter ‘*f*’ followed by a picture of a *flamingo*). For each presented stimulus, subjects viewed a fixation cross for 500 ms, a single-letter probe for 250ms, the object image (or word) for 500 ms, followed by 2–7 s of jitter. On Day 2, subjects completed a conceptual memory test in the scanner where they were presented with word labels for old objects that were previously seen (*n* = 300) and new objects that were not previously seen (*n* = 100). Subjects rated their confidence (1 = “definitely new”, 2 = “probably new”, 3 = “probably old”, and 4 = “definitely old”) for whether they had seen the object the word referred to on Day 1.

#### MRI Acquisition

Scans were performed on a GE MR 750 3T scanner. Coplanar functional images were acquired using an inverse spiral sequence: 37 axial slices, 64×64 matrix, in-plane resolution 4×4mm^2^, 3.8mm slice thickness, flip angle = 77, TR = 2000ms, TE 31ms, FOV = 24mm^2^. Anatomical images were acquired using a 3D T1-weighted echo-planar sequence (68 slices, 256×256 matrix, in-plane resolution 2×2mm^2^, 1.9mm slice thickness, TR=12 ms, TE=5mg, FOV=24cm). Encoding was completed in two separate runs and retrieval was completed in four separate runs.

Data preprocessing was performed using SPM12 and custom MATLAB scripts. Functional images were realigned to the first image of the first run using rigid-body transformation with six motion parameters. Functional images were then corrected for slice acquisition time (reference slice = first slice) and linear signal drift, temporally smoothed using a high-pass filter of 190s, segmented into different tissue types (gray matter, white matter, and cerebrospinal fluid), and normalized to the MNI152 standard space. Automatic ICA-denoising was applied to remove artifacts due to motion or susceptibility artifacts and the effects of head motion, button presses, white matter signals, and cerebrospinal fluid signals were included as 1^st^-level covariates of no interest. We used Least-Squares Separate, a technique designed for signal estimating in event-related designs (Mumford et al., 2012, 2014; Rissman et al., 2004). For each gray matter voxel, the activity estimate on each encoding trial was obtained by constructing a first-level general linear model with a one regressor for each trial and all other trials collapsed into a second regressor. This process was repeated for each trial, resulting in one beta estimate per trial.

#### Neuroimaging Analysis

For each subject, beta images from each single-trial model were used to estimate activity associated with each object, by averaging all voxel values within a given region of interest (ROI). Forty ROIs, defined using the Brainnetome atlas (Fan et al., 2016), were used in the present analysis: anterior temporal lobe, fusiform gyrus, hippocampus, inferior frontal gyrus, inferior parietal lobule, inferior temporal gyrus, lateral occipital cortex, medioventral occipital cortex, middle temporal gyrus, precuneus, perirhinal cortex, parahippocampal cortex, parahippocampal gyrus, postcentral gyrus, precentral gyrus, posterior superior temporal sulcus, rhinal cortex, retrosplenial cortex, superior temporal gyrus, supramarginal gyrus. They were chosen because previous work has shown them to be crucial for semantic classification (Usami et al., 2022, Clark et al., 2014). We used unilateral ROIs for all 20 regions (making a total of 40). For each participant, trial-level activation level was computed as the average of activity estimates (betas) for a given object stimulus on a single trial across voxels within an ROI mask using a general linear model. Brain images were visualized using the FSLeyes toolbox (fsl.fmrib.ox.ac.uk/fsl/fslwiki/FSLeyes).

#### Nonnegative Matrix Factorization

A principal outcome of Experiment 1 was a Concept × Feature frequency matrix describing 5520 unique features for each of 995 real-world object. Each individual cell indicates the frequency with which a given feature was provided for a given object. Two individual feature matrices were used in the analysis based on McRae feature norms: encyclopedic and visual (McRae et al., 2005). Additional feature categories such as taxonomic features were not isolated for additional analysis though were still present in the original feature matrix.

In order to determine a robust set of semantic factors from the feature norms we obtained in Experiment 1, we used Nonnegative Matrix Factorization (NMF) (Paatero & Tapper, 1994). NMF takes an input matrix V and factorizes into two matrices W and H that, when multiplied, give a lower-rank approximation of the original matrix. NMF is preferable to other commonly used dimensionality reduction techniques such as Principal Components Analysis (PCA) and Factor Analysis (FA) when a dataset is sparse and linearity cannot be assumed (Paatero & Tapper, 1994). Furthermore, nonnegativity tends to produce more interpretable dimensions when compared with other dimensionality reduction methods that don’t use such a constraint (Roads & Love, 2024). While many object features are shared (e.g., “has legs”, “is made of metal”), most features are unique to a small minority of objects, and as such the Object x Feature matrix is quite sparse (5,450,054 zero elements out of a total 5,492,400 elements, for a sparsity ratio of 0.9923). NMF is especially appropriate for such sparse matrices, because it derives matrices W (feature weights) and H (coefficients), both non-negative factors, which together constitute a lower-rank approximation of a given matrix (see Figure 3).

**Figure 1.**
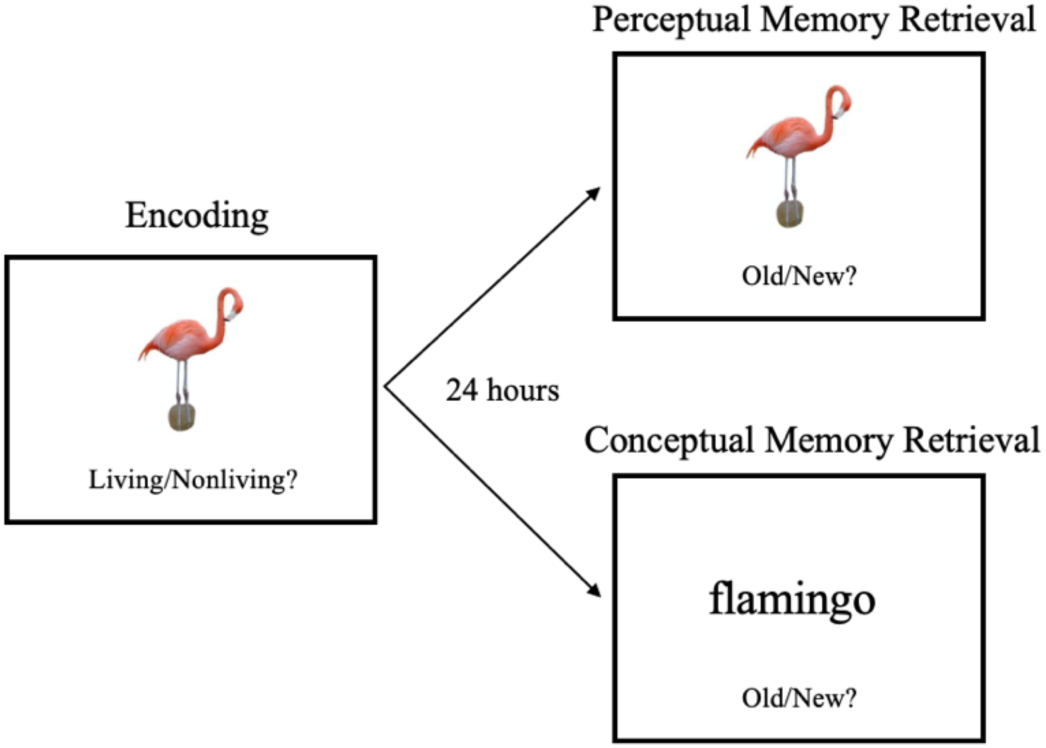
Memorability Task. Participants view a series of images and rate whether the object is living or nonliving. After a 24-hour delay, the participants were divided into two groups and completed either the perceptual or conceptual recognition memory task. They determined whether a given image or word was seen on the previous day (i.e., old) or was not (i.e., new).

**Figure 2.**
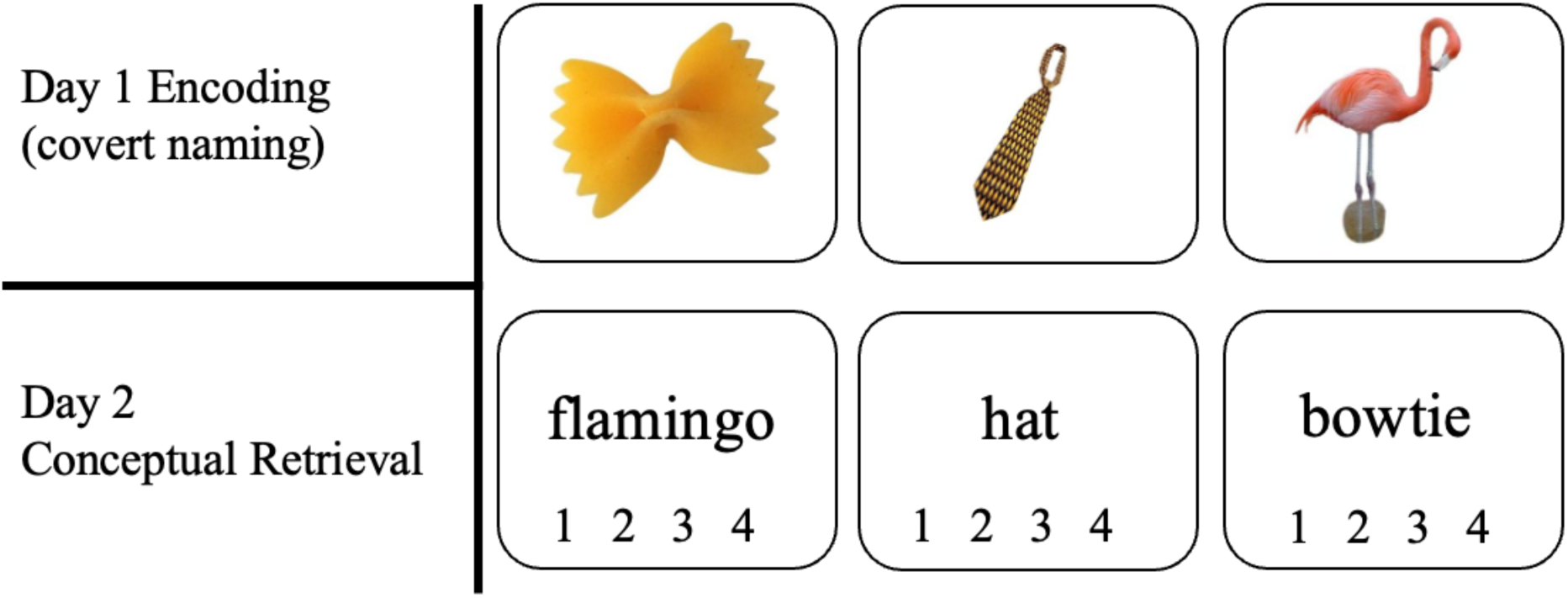
Memory Task for Experiment 3. Participants viewed images during an MRI scan and covertly identified that image, such as *bowtie pasta* or *flamingo*. On the second day, participants were scanned while viewing words in a recognition memory task. If the image was seen on Day 1, they would rate 3 or 4 for probably or definitely old. If the image was not seen on Day 1, they would rate 1 or 2 for definitely or probably new.

**Figure 3.**
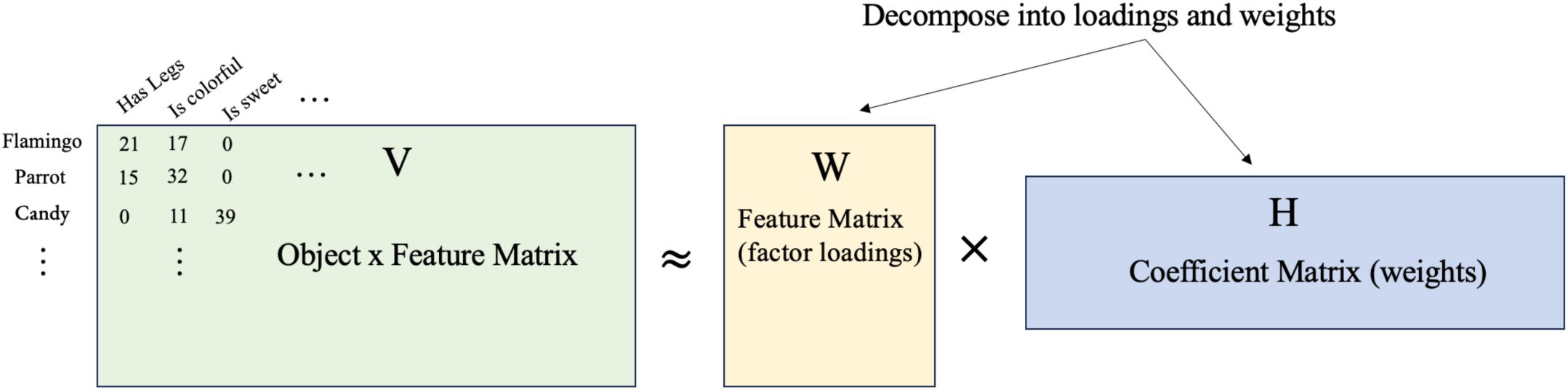
Nonnegative Matrix Factorization of Feature Norms. Conceptual description of the nonnegative matrix factorization procedure applied to the feature norm matrix. The V matrix consisted of object stimuli and the number of times a given feature was provided by participants in the feature norming study (Experiment 1). NMF decomposes this matrix into a feature matrix W and coefficient matrix H, which when recombined give an approximation of the original V matrix.

#### Applying NMF

After completing the thresholding procedure (see Supplementary Figure 1), we applied a standard implementation of NMF from MATLAB (Mathworks, Inc.) using the *nnmf* function. We chose the standard alternative least squares algorithm (Paatero, 1997). The use of NMF in cognitive neuroscience has shown mixed utility. While the method shows success for structural MRI (Patel et al., 2020; Sotiras et al., 2015), it has been less effective for functional MRI (Xie et al., 2017). NMF is more widely used for psychology and computing applications involving language (Hassani et al., 2021; Mangin et al., 2015) and emotion (Calvo & Mac Kim, 2013), suggesting that it is appropriately chosen to model object features and semantics. We applied several methods for determining the optimal number of factors to extract from the W matrix, identifying 20 to use in subsequent analyses (see **Supplementary** Figure 2). It is important to note that while NMF does not rank factors in order of explanatory importance, we discovered that approximately the first five factors exhibited greater explanatory power. We assessed this by reconstructing the original V matrix from the component W and H matrices, excluding one factor at a time.

#### Identifying Cortical Memorability Effects

In order to model the relationship between brain activity, normative memorability, and memory performance for the subjects in the neuroimaging study, we first calculated corrected recognition (Hit rate – False Alarm rate) for all trials for each of the 19 subjects for Conceptual and Perceptual Memory (i.e., individual subject memory performance from Experiment 3, whereas normative memorability came from a separate group of subjects in Experiment 2). Due to the need to account for subject-wise variability, we fit a linear mixed-effects model using the “lme4” package (Bates et al., 2015) in R (version 4.3.2). We submitted trial-level brain activity to a mixed-effects model with the fixed effects of conceptual corrected recognition, perceptual corrected recognition, and object with a random effect of subject. Task performance between conceptual and perceptual memory tasks in Experiment 3 (not shown in figure) were positively correlated, so perceptual corrected recognition was included as a regressor to properly estimate the effects of memory regardless of the performance of the neuroimaging subjects. We then used the estimate for each object as the adjusted brain activity for that trial. We then modeled the adjusted brain activity with a linear model with a fixed effect of normative memorability, yielding a subset of significant ROIs from our list of ROIs. After taking these steps to account for object-wise performance across subjects, analyses were conducted using only subsequently remembered trials because subsequently forgotten trials may be evidence of inattention towards that particular image, potentially resulting in a lack of representation of the image.

#### Mediation Analysis

In order to investigate whether semantics mediate the relationship between memorability and brain activity, we performed a mediation analysis using the *lavaan* package for Latent Variable Analysis in R (Rosseel et al., 2024). A multilevel mediation model was run to investigate conceptual memorability, five semantic factors, and all ROIs. Subsequent models were run to investigate the mediating influence of three factor types (encyclopedic, visual, and functional factors). The multilevel mediation model considered each mediator in parallel, such that the relationship between memorability and each semantic factor is a separate *a* path, the relationship between semantic factors and activity in each ROI is given by a separate *b* path, and the relationship between memorability and brain activity in each ROI is the *c* path (Figure 5). A significant *a × b* interaction showed that semantic factors were significant mediators of the relationship between memorability and activity. The *a × b* interaction effects were tested with bootstrap confidence intervals. An additional benefit of multilevel mediation is that it affords control over multiple repeated factors (e.g., each stimulus). The variables at Level 1 included trial-wise activity in a given ROI for each subject and the intercepts were added at Level 2, along with object-wise memorability and the object-wise loading on each semantic factor. In this way we can model the estimated brain activity across participants for each stimulus and complete an object-wise mediation analysis. Mediation models were fit using the Structural Equation Modeling function in R (Fox et al., 2022), after which Huber-White standard errors were calculated and parameter estimates extracted.

## RESULTS

### Semantic Dimensions

Our NMF procedure identified five factors capable of reconstructing the concept-feature frequency matrices for encyclopedic and visual features. Each factor gave a unique pattern of loadings for different objects (W matrix) and features (H matrix) (Figure 4). For example, consider the W and H matrices derived from the matrix of encyclopedic features. The H matrix gives coefficients for each feature and the W matrix gives loadings for each object. For the encyclopedic factors, Factor 1 was associated with the following features (sorted in descending order by their coefficients): “is useful,” “is helpful,” and “is portable” (Figure 6). The coefficients indicate how important that feature is for the factor. By focusing on the largest loading values, we can approximate an identity for that factor. In this case, *usefulness* is a defining feature of Factor 1. Each object is associated with a loading on each factor in the W matrix. The top objects for Factor 1 include *bucket*, *trolley*, and *camel*. While these objects do not belong to one group based on a standard hierarchical categorization scheme, they clearly reflect the concept of *usefulness* in their respective contexts. Usefulness is an important feature of tools, as well as animals and other objects which are characterized by their utility. In contrast, Factor 2, which has top features that include: “does fly,” “does eat fish,” and “does lay eggs” more clearly recapitulates a traditional category, reflecting the general category of *animacy*.

**Figure 4.**
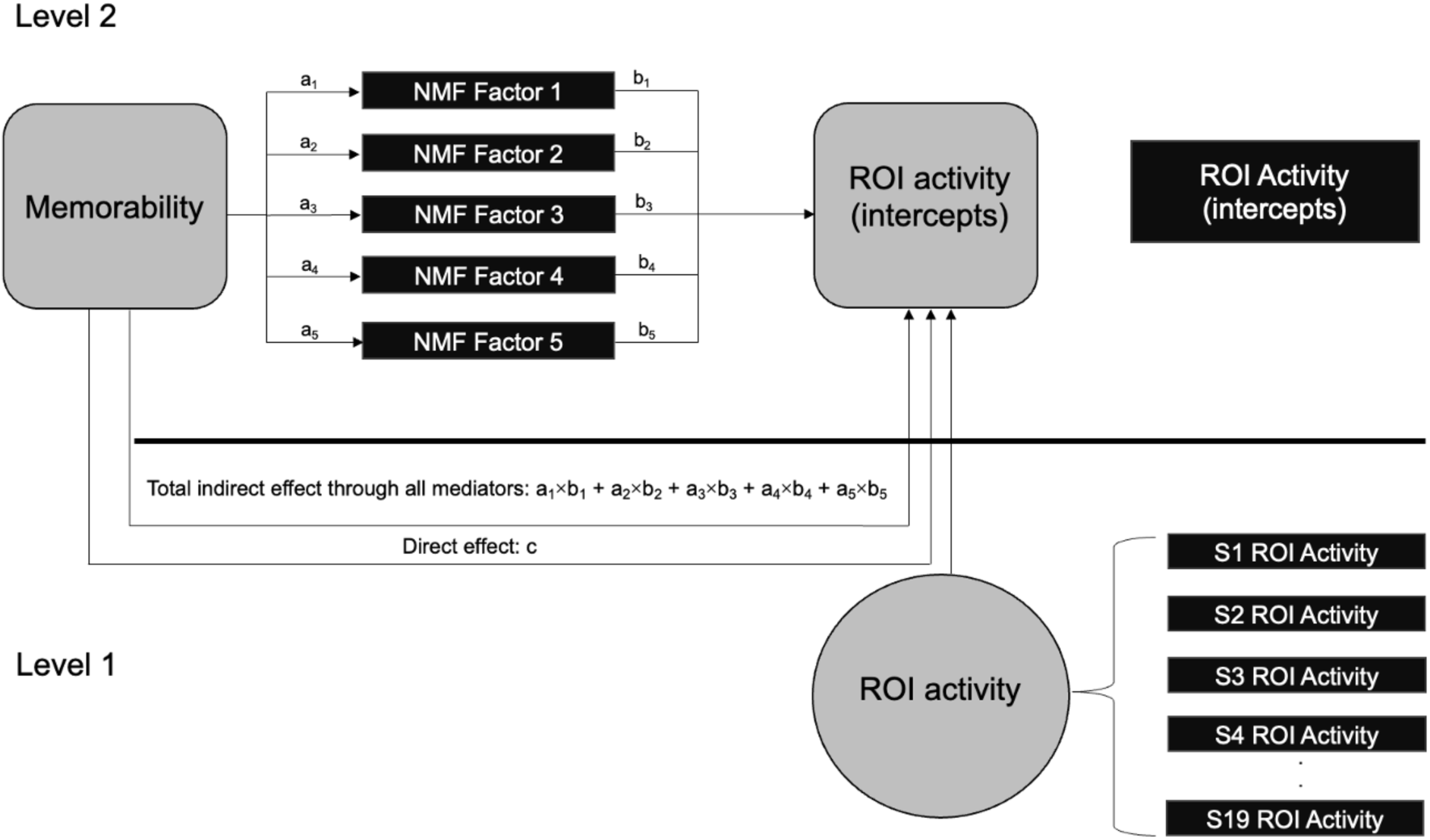
Multilevel mediation with five mediators. In this model, a 2-2-1, the outcome variable is at level 1 because it refers to the brain activity of individual subjects, whereas the predictor and mediator at level 2 because these properties of the stimuli remain constant for all participants. In this way, the mediation analysis takes place at level 2, using the intercepts to capture the brain activity across participants for each stimulus. Such an object-wise model allows for the examination of how stimulus characteristics influence neural responses while accounting for both between-stimulus and between-subject variability.

**Figure 5.**
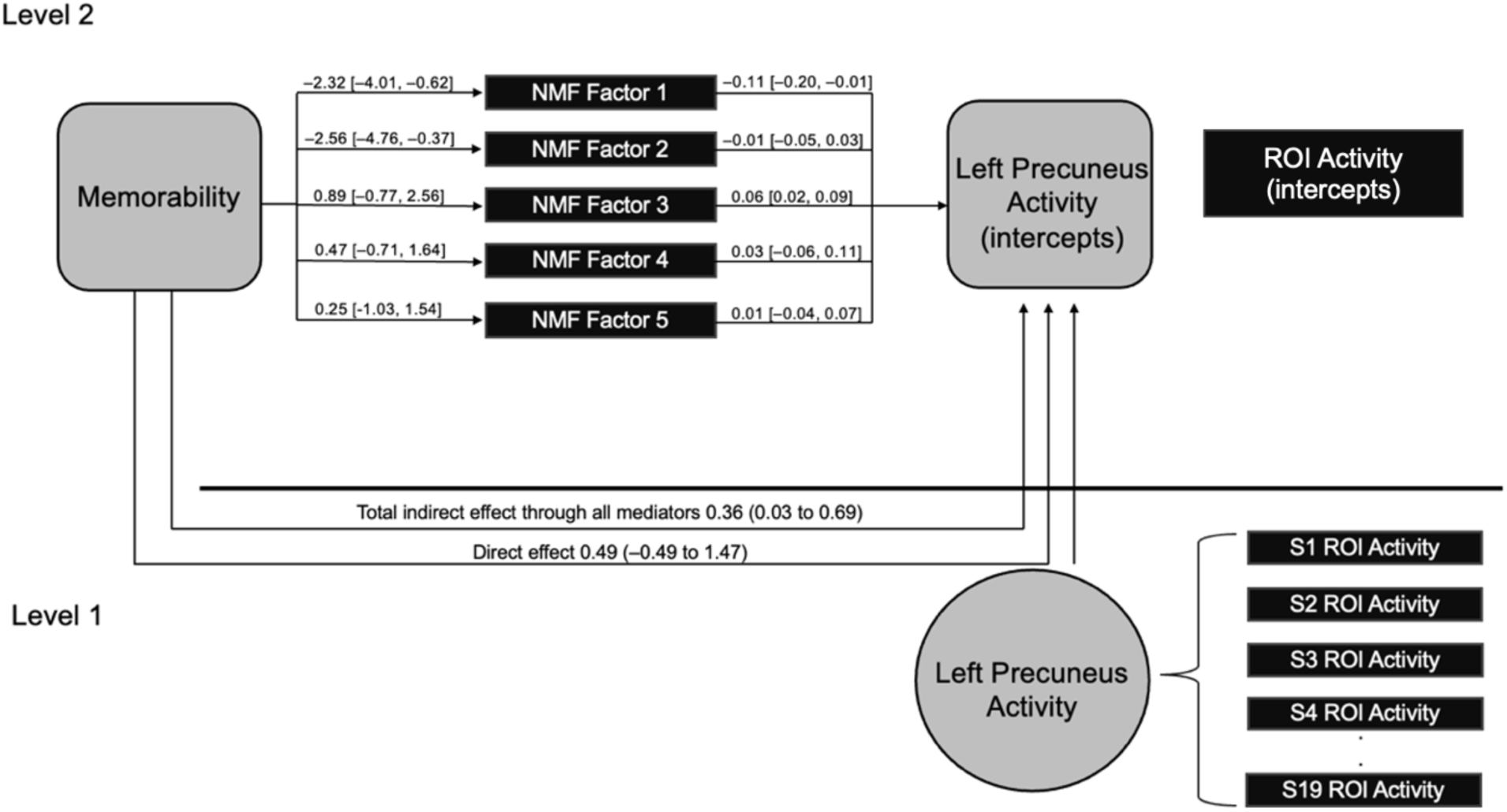
Sample multilevel mediation in one ROI. Here is a sample mediation model where the outcome variability is trial-wise activity in left precuneus. The stimuli presented at each trial determine the predictor (i.e., memorability) and mediators (i.e., the semantic factors). We estimate each path in the mediation model and can calculate the direct effect of the predictor on the outcome variable and the indirect effect through the mediators.

**Figure 6.**
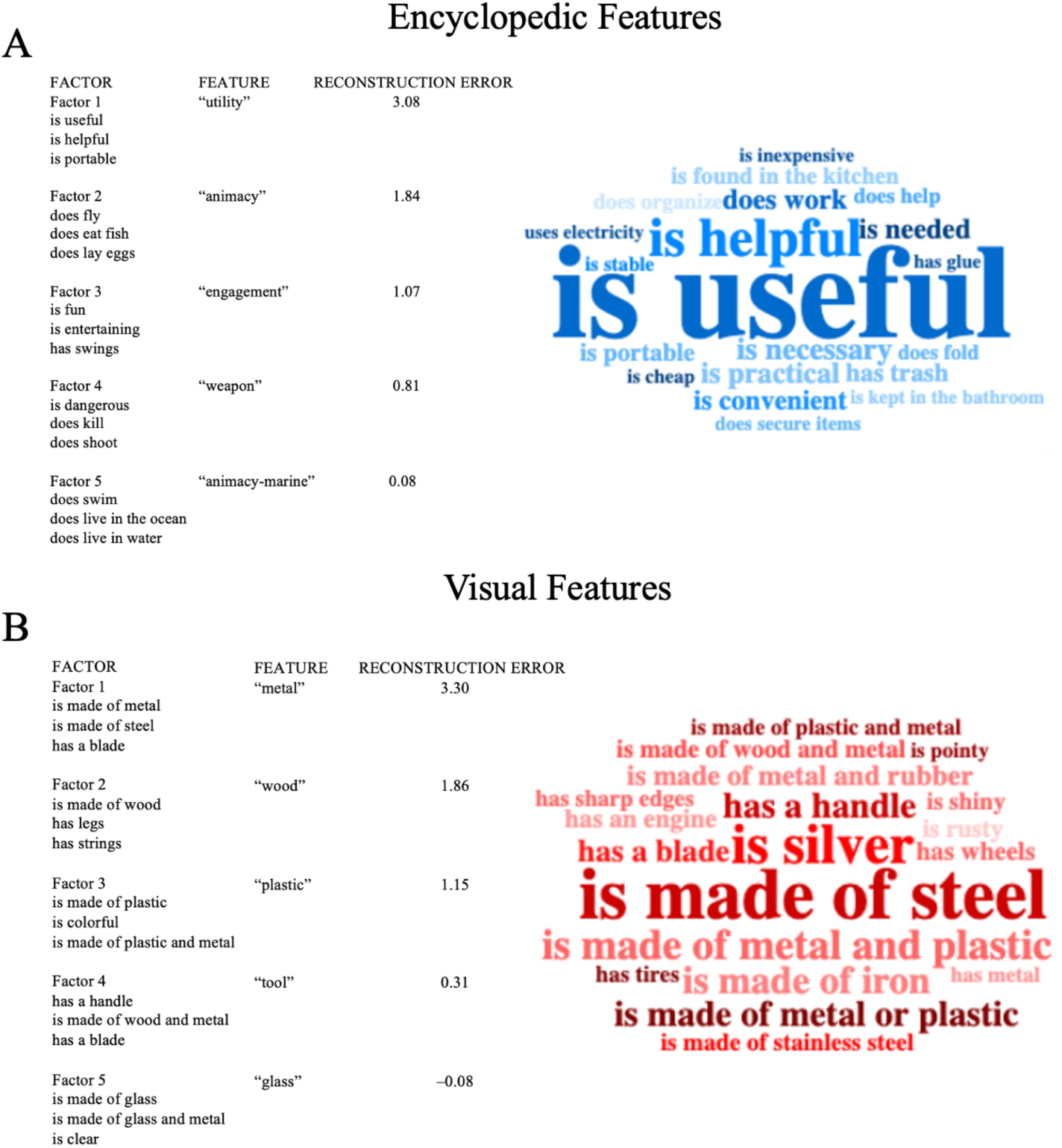
Semantic factor features. Top factors based on NMF for encyclopedic (A) and visual (B) features, with the top three features contributing to the factor listed. The reconstruction error is a proxy for the factors importance because it quantifies how much it contributes to the overall reconstruction of the original concept-feature frequency matrix (a higher reconstruction error means a given feature is more important). To visualize which features contribute to the top encyclopedic and top visual features, word clouds representing the relative importance of features are presented.

**Figure 7.**
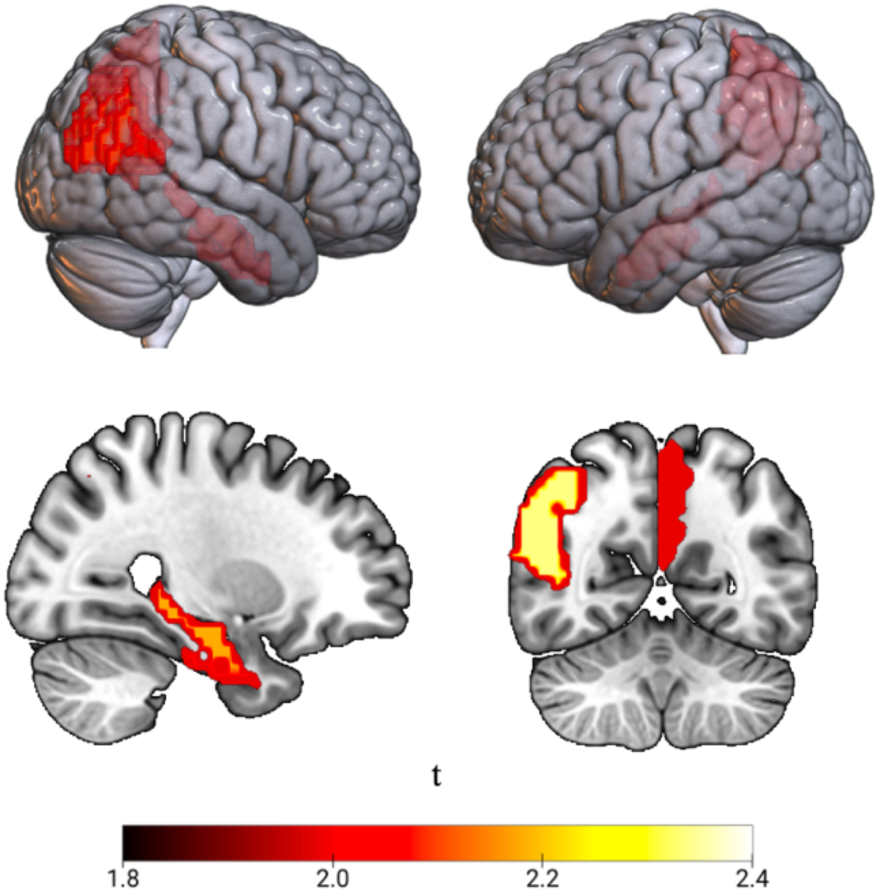
Memorability-related regions. Using a general linear model, we examined the relationship between stimulus memorability and univariate brain activity in our ROIs for hit trials only. Five regions showed significant modulation of trial-wise activity in response to the memorability of the stimuli presented during those trials. The figure displays a t-map which is colored based on the corresponding t-value from the analysis. Degrees of freedom reported as (model df, residual df).

Factor 3, characterized by top features like “is fun” and “is entertaining” has top-loading objects such as *swing set*, *saxophone*, and *pickle*, indicating that the semantic attribute of “fun” can apply to toys, musical instruments, and food, among others. Therefore, each factor captures semantic features common to objects from a wide range of categories.

The top visual factors offer dimensions with a more straightforward taxonomy, often clearly based on the material of objects, such as being made of metal (Factor 1), wood (Factor 2), plastic (Factor 3), or glass (Factor 5), as well as reflecting the general category of tools (Factor 4).

Interestingly, NMF applied to the original feature matrix (which includes all feature types) identifies material-based features (e.g. made of metal or wood), as well as object categories such as *animal* or *tool*.

### Memorability-related regions

Our first analysis of fMRI data focused on identifying cortical regions that show a significant sensitivity to variability in object memorability (measured by corrected recognition). Using object-wise conceptual and perceptual normative memorability from Experiment 2 and the adjusted trial-wise brain activity from Experiment 3, we performed linear modeling to determine which ROIs showed a significant relationship between brain activity and object memorability (**Table 1**). Of our original 40 ROIs, five showed a significant effect, where the more memorable the object the more brain activity: the right hippocampus (β = 0.38, *t*(2) = 2.18, *p* = 0.03), left precuneus (β = 1.1, *t*(2) = 1.99, *p* = 0.048), right rhinal cortex (β *=* 0.32*, t*(2) = 2.07, *p* = 0.04), right inferior parietal lobule (β *=* 0.76*, t*(2) = 2.34, *p* = 0.02) and right superior temporal gyrus (β = 0.58, *t*(2) = 1.98, *p* = 0.049). These results are consistent with other reports demonstrating object-level variance in memorability in ventral temporal cortex (Bainbridge & Rissman, 2018), as well as with the more general role of this system in contributing to successful object memory (Tyler et al., 2013).

**Table 1.** Effects of memorability on cortical activity

| Region | Beta | SE | df | t-value | p-value |
| --- | --- | --- | --- | --- | --- |
| Hippocampus R | 0.38 | 0.18 | (2, 240) | 2.18 | 0.0302 |
| Precuneus L | 1.10 | 0.55 | (2, 240) | 1.99 | 0.0482 |
| Rhinal Cortex R | 0.32 | 0.15 | (2, 240) | 2.07 | 0.0397 |
| Inferior Parietal Lobule R | 0.76 | 0.33 | (2, 240) | 2.34 | 0.0203 |
| Superior Temporal Gyrus R | 0.58 | 0.30 | (2, 240) | 1.98 | 0.0492 |

**Table 1.**
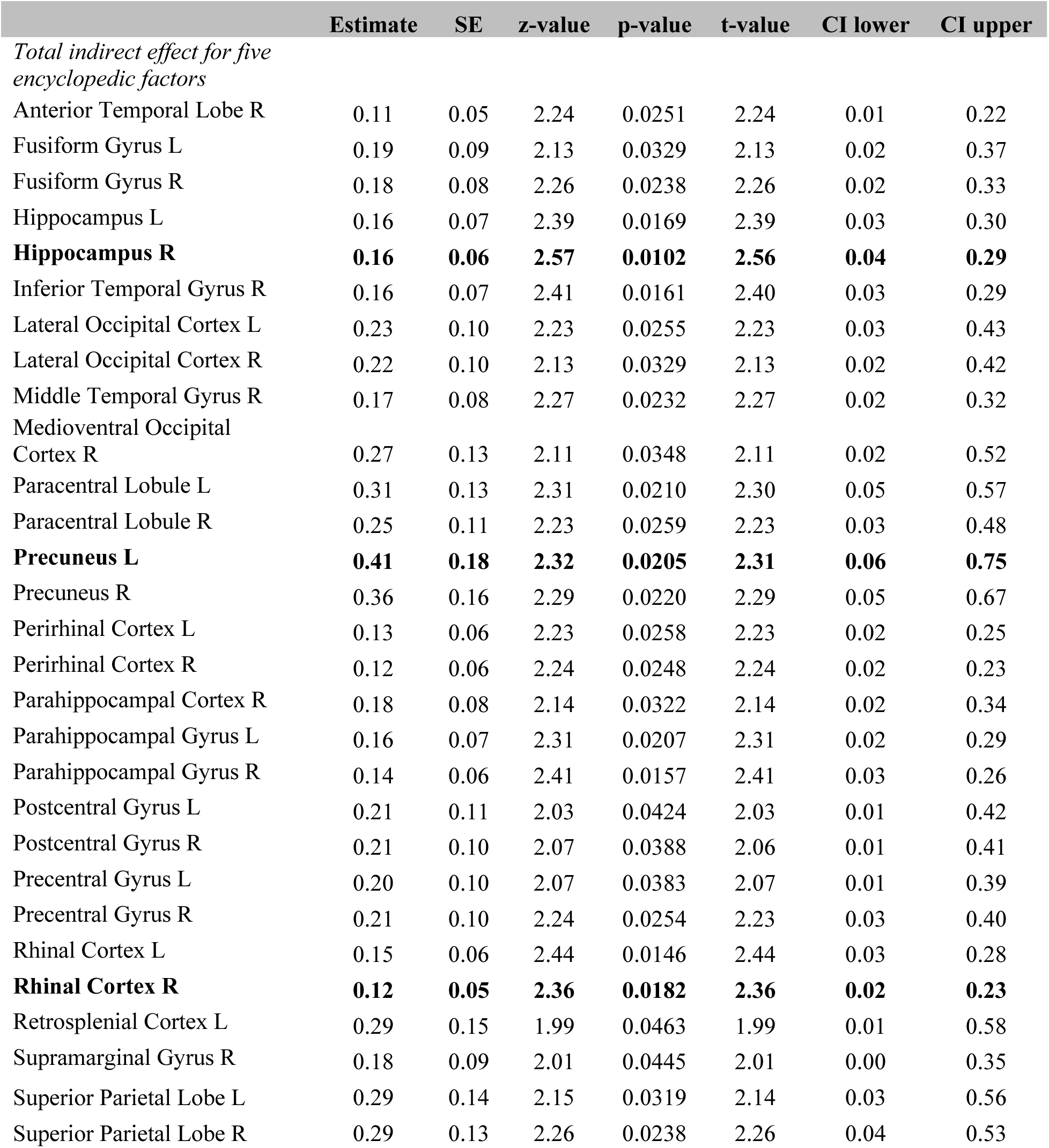
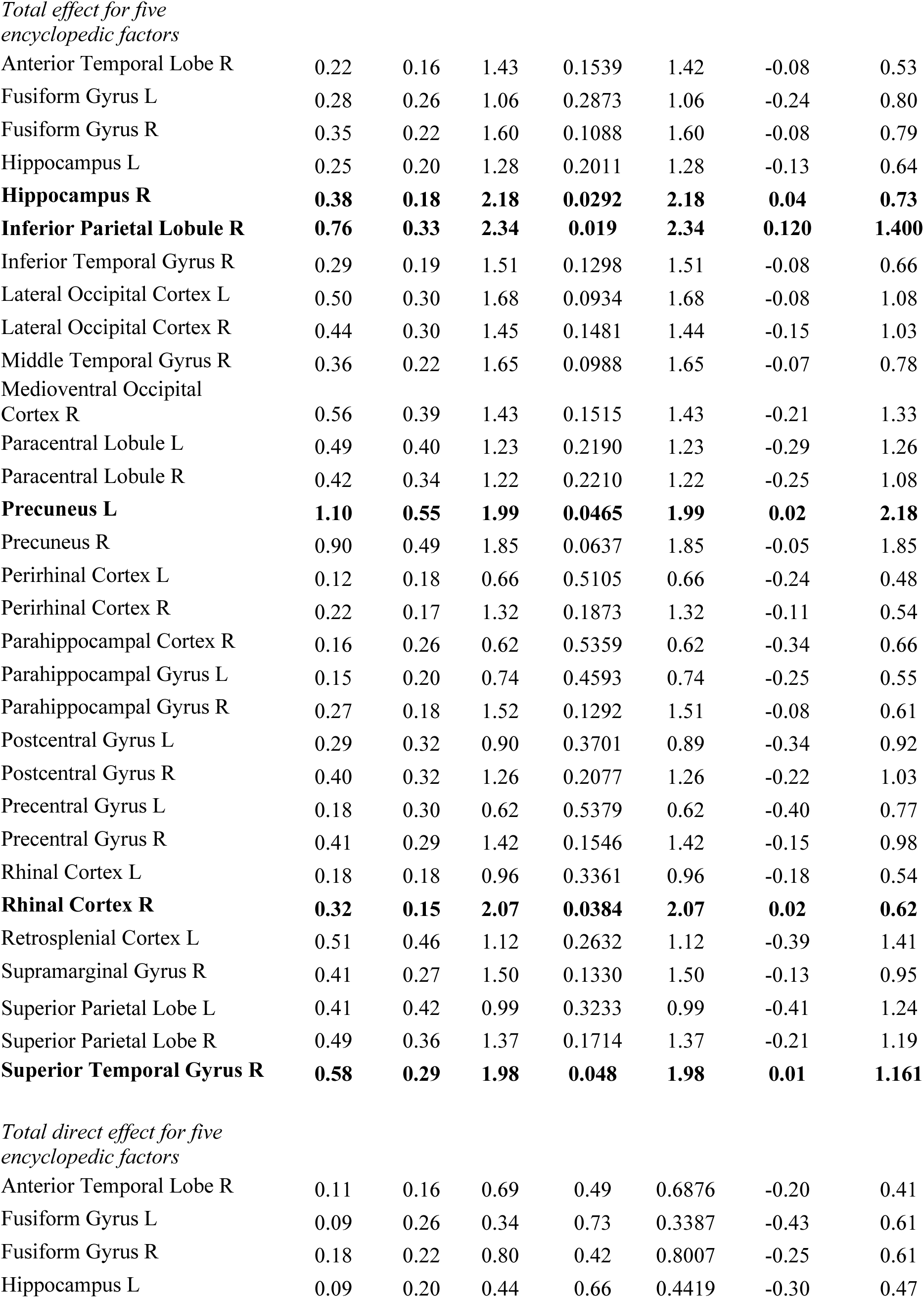
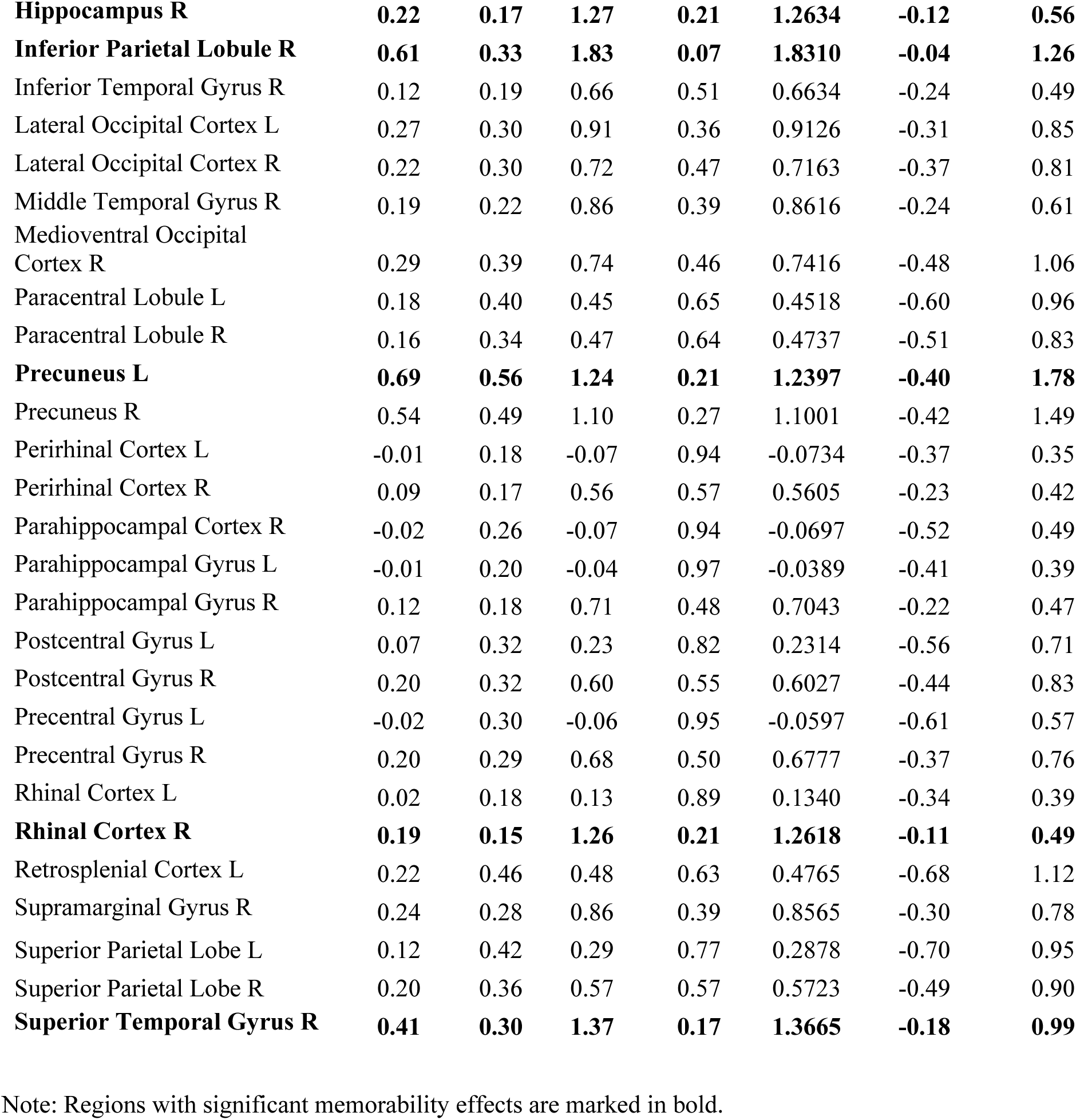
Mediation results for encyclopedic factors

|  | Estimate | SE | z-value | p-value | t-value | CI lower | CI upper |
| --- | --- | --- | --- | --- | --- | --- | --- |
| *Total indirect effect for five encyclopedic factors* |  |  |  |  |  |  |  |
| Anterior Temporal Lobe R | 0.11 | 0.05 | 2.24 | 0.0251 | 2.24 | 0.01 | 0.22 |
| Fusiform Gyrus L | 0.19 | 0.09 | 2.13 | 0.0329 | 2.13 | 0.02 | 0.37 |
| Fusiform Gyrus R | 0.18 | 0.08 | 2.26 | 0.0238 | 2.26 | 0.02 | 0.33 |
| Hippocampus L | 0.16 | 0.07 | 2.39 | 0.0169 | 2.39 | 0.03 | 0.30 |
| **Hippocampus R** | **0.16** | **0.06** | **2.57** | **0.0102** | **2.56** | **0.04** | **0.29** |
| Inferior Temporal Gyrus R | 0.16 | 0.07 | 2.41 | 0.0161 | 2.40 | 0.03 | 0.29 |
| Lateral Occipital Cortex L | 0.23 | 0.10 | 2.23 | 0.0255 | 2.23 | 0.03 | 0.43 |
| Lateral Occipital Cortex R | 0.22 | 0.10 | 2.13 | 0.0329 | 2.13 | 0.02 | 0.42 |
| Middle Temporal Gyrus R | 0.17 | 0.08 | 2.27 | 0.0232 | 2.27 | 0.02 | 0.32 |
| Medioventral Occipital Cortex R | 0.27 | 0.13 | 2.11 | 0.0348 | 2.11 | 0.02 | 0.52 |
| Paracentral Lobule L | 0.31 | 0.13 | 2.31 | 0.0210 | 2.30 | 0.05 | 0.57 |
| Paracentral Lobule R | 0.25 | 0.11 | 2.23 | 0.0259 | 2.23 | 0.03 | 0.48 |
| **Precuneus L** | **0.41** | **0.18** | **2.32** | **0.0205** | **2.31** | **0.06** | **0.75** |
| Precuneus R | 0.36 | 0.16 | 2.29 | 0.0220 | 2.29 | 0.05 | 0.67 |
| Perirhinal Cortex L | 0.13 | 0.06 | 2.23 | 0.0258 | 2.23 | 0.02 | 0.25 |
| Perirhinal Cortex R | 0.12 | 0.06 | 2.24 | 0.0248 | 2.24 | 0.02 | 0.23 |
| Parahippocampal Cortex R | 0.18 | 0.08 | 2.14 | 0.0322 | 2.14 | 0.02 | 0.34 |
| Parahippocampal Gyrus L | 0.16 | 0.07 | 2.31 | 0.0207 | 2.31 | 0.02 | 0.29 |
| Parahippocampal Gyrus R | 0.14 | 0.06 | 2.41 | 0.0157 | 2.41 | 0.03 | 0.26 |
| Postcentral Gyrus L | 0.21 | 0.11 | 2.03 | 0.0424 | 2.03 | 0.01 | 0.42 |
| Postcentral Gyrus R | 0.21 | 0.10 | 2.07 | 0.0388 | 2.06 | 0.01 | 0.41 |
| Precentral Gyrus L | 0.20 | 0.10 | 2.07 | 0.0383 | 2.07 | 0.01 | 0.39 |
| Precentral Gyrus R | 0.21 | 0.10 | 2.24 | 0.0254 | 2.23 | 0.03 | 0.40 |
| Rhinal Cortex L | 0.15 | 0.06 | 2.44 | 0.0146 | 2.44 | 0.03 | 0.28 |
| **Rhinal Cortex R** | **0.12** | **0.05** | **2.36** | **0.0182** | **2.36** | **0.02** | **0.23** |
| Retrosplenial Cortex L | 0.29 | 0.15 | 1.99 | 0.0463 | 1.99 | 0.01 | 0.58 |
| Supramarginal Gyrus R | 0.18 | 0.09 | 2.01 | 0.0445 | 2.01 | 0.00 | 0.35 |
| Superior Parietal Lobe L | 0.29 | 0.14 | 2.15 | 0.0319 | 2.14 | 0.03 | 0.56 |
| Superior Parietal Lobe R | 0.29 | 0.13 | 2.26 | 0.0238 | 2.26 | 0.04 | 0.53 |
| Anterior Temporal Lobe R | 0.22 | 0.16 | 1.43 | 0.1539 | 1.42 | -0.08 | 0.53 |
| Fusiform Gyrus L | 0.28 | 0.26 | 1.06 | 0.2873 | 1.06 | -0.24 | 0.80 |
| Fusiform Gyrus R | 0.35 | 0.22 | 1.60 | 0.1088 | 1.60 | -0.08 | 0.79 |
| Hippocampus L | 0.25 | 0.20 | 1.28 | 0.2011 | 1.28 | -0.13 | 0.64 |
| **Hippocampus R** | **0.38** | **0.18** | **2.18** | **0.0292** | **2.18** | **0.04** | **0.73** |
| **Inferior Parietal Lobule R** | **0.76** | **0.33** | **2.34** | **0.019** | **2.34** | **0.120** | **1.400** |
| Inferior Temporal Gyrus R | 0.29 | 0.19 | 1.51 | 0.1298 | 1.51 | -0.08 | 0.66 |
| Lateral Occipital Cortex L | 0.50 | 0.30 | 1.68 | 0.0934 | 1.68 | -0.08 | 1.08 |
| Lateral Occipital Cortex R | 0.44 | 0.30 | 1.45 | 0.1481 | 1.44 | -0.15 | 1.03 |
| Middle Temporal Gyrus R | 0.36 | 0.22 | 1.65 | 0.0988 | 1.65 | -0.07 | 0.78 |
| Medioventral Occipital<br>Cortex R | 0.56 | 0.39 | 1.43 | 0.1515 | 1.43 | -0.21 | 1.33 |
| Paracentral Lobule L | 0.49 | 0.40 | 1.23 | 0.2190 | 1.23 | -0.29 | 1.26 |
| Paracentral Lobule R | 0.42 | 0.34 | 1.22 | 0.2210 | 1.22 | -0.25 | 1.08 |
| **Precuneus L** | **1.10** | **0.55** | **1.99** | **0.0465** | **1.99** | **0.02** | **2.18** |
| Precuneus R | 0.90 | 0.49 | 1.85 | 0.0637 | 1.85 | -0.05 | 1.85 |
| Perirhinal Cortex L | 0.12 | 0.18 | 0.66 | 0.5105 | 0.66 | -0.24 | 0.48 |
| Perirhinal Cortex R | 0.22 | 0.17 | 1.32 | 0.1873 | 1.32 | -0.11 | 0.54 |
| Parahippocampal Cortex R | 0.16 | 0.26 | 0.62 | 0.5359 | 0.62 | -0.34 | 0.66 |
| Parahippocampal Gyrus L | 0.15 | 0.20 | 0.74 | 0.4593 | 0.74 | -0.25 | 0.55 |
| Parahippocampal Gyrus R | 0.27 | 0.18 | 1.52 | 0.1292 | 1.51 | -0.08 | 0.61 |
| Postcentral Gyrus L | 0.29 | 0.32 | 0.90 | 0.3701 | 0.89 | -0.34 | 0.92 |
| Postcentral Gyrus R | 0.40 | 0.32 | 1.26 | 0.2077 | 1.26 | -0.22 | 1.03 |
| Precentral Gyrus L | 0.18 | 0.30 | 0.62 | 0.5379 | 0.62 | -0.40 | 0.77 |
| Precentral Gyrus R | 0.41 | 0.29 | 1.42 | 0.1546 | 1.42 | -0.15 | 0.98 |
| Rhinal Cortex L | 0.18 | 0.18 | 0.96 | 0.3361 | 0.96 | -0.18 | 0.54 |
| **Rhinal Cortex R** | **0.32** | **0.15** | **2.07** | **0.0384** | **2.07** | **0.02** | **0.62** |
| Retrosplenial Cortex L | 0.51 | 0.46 | 1.12 | 0.2632 | 1.12 | -0.39 | 1.41 |
| Supramarginal Gyrus R | 0.41 | 0.27 | 1.50 | 0.1330 | 1.50 | -0.13 | 0.95 |
| Superior Parietal Lobe L | 0.41 | 0.42 | 0.99 | 0.3233 | 0.99 | -0.41 | 1.24 |
| Superior Parietal Lobe R | 0.49 | 0.36 | 1.37 | 0.1714 | 1.37 | -0.21 | 1.19 |
| **Superior Temporal Gyrus R** | **0.58** | **0.29** | **1.98** | **0.048** | **1.98** | **0.01** | **1.161** |
| Anterior Temporal Lobe R | 0.11 | 0.16 | 0.69 | 0.49 | 0.6876 | -0.20 | 0.41 |
| Fusiform Gyrus L | 0.09 | 0.26 | 0.34 | 0.73 | 0.3387 | -0.43 | 0.61 |
| Fusiform Gyrus R | 0.18 | 0.22 | 0.80 | 0.42 | 0.8007 | -0.25 | 0.61 |
| Hippocampus L | 0.09 | 0.20 | 0.44 | 0.66 | 0.4419 | -0.30 | 0.47 |
| **Hippocampus R** | **0.22** | **0.17** | **1.27** | **0.21** | **1.2634** | **-0.12** | **0.56** |
| **Inferior Parietal Lobule R** | **0.61** | **0.33** | **1.83** | **0.07** | **1.8310** | **-0.04** | **1.26** |
| Inferior Temporal Gyrus R | 0.12 | 0.19 | 0.66 | 0.51 | 0.6634 | -0.24 | 0.49 |
| Lateral Occipital Cortex L | 0.27 | 0.30 | 0.91 | 0.36 | 0.9126 | -0.31 | 0.85 |
| Lateral Occipital Cortex R | 0.22 | 0.30 | 0.72 | 0.47 | 0.7163 | -0.37 | 0.81 |
| Middle Temporal Gyrus R | 0.19 | 0.22 | 0.86 | 0.39 | 0.8616 | -0.24 | 0.61 |
| Medioventral Occipital Cortex R | 0.29 | 0.39 | 0.74 | 0.46 | 0.7416 | -0.48 | 1.06 |
| Paracentral Lobule L | 0.18 | 0.40 | 0.45 | 0.65 | 0.4518 | -0.60 | 0.96 |
| Paracentral Lobule R | 0.16 | 0.34 | 0.47 | 0.64 | 0.4737 | -0.51 | 0.83 |
| **Precuneus L** | **0.69** | **0.56** | **1.24** | **0.21** | **1.2397** | **-0.40** | **1.78** |
| Precuneus R | 0.54 | 0.49 | 1.10 | 0.27 | 1.1001 | -0.42 | 1.49 |
| Perirhinal Cortex L | -0.01 | 0.18 | -0.07 | 0.94 | -0.0734 | -0.37 | 0.35 |
| Perirhinal Cortex R | 0.09 | 0.17 | 0.56 | 0.57 | 0.5605 | -0.23 | 0.42 |
| Parahippocampal Cortex R | -0.02 | 0.26 | -0.07 | 0.94 | -0.0697 | -0.52 | 0.49 |
| Parahippocampal Gyrus L | -0.01 | 0.20 | -0.04 | 0.97 | -0.0389 | -0.41 | 0.39 |
| Parahippocampal Gyrus R | 0.12 | 0.18 | 0.71 | 0.48 | 0.7043 | -0.22 | 0.47 |
| Postcentral Gyrus L | 0.07 | 0.32 | 0.23 | 0.82 | 0.2314 | -0.56 | 0.71 |
| Postcentral Gyrus R | 0.20 | 0.32 | 0.60 | 0.55 | 0.6027 | -0.44 | 0.83 |
| Precentral Gyrus L | -0.02 | 0.30 | -0.06 | 0.95 | -0.0597 | -0.61 | 0.57 |
| Precentral Gyrus R | 0.20 | 0.29 | 0.68 | 0.50 | 0.6777 | -0.37 | 0.76 |
| Rhinal Cortex L | 0.02 | 0.18 | 0.13 | 0.89 | 0.1340 | -0.34 | 0.39 |
| **Rhinal Cortex R** | **0.19** | **0.15** | **1.26** | **0.21** | **1.2618** | **-0.11** | **0.49** |
| Retrosplenial Cortex L | 0.22 | 0.46 | 0.48 | 0.63 | 0.4765 | -0.68 | 1.12 |
| Supramarginal Gyrus R | 0.24 | 0.28 | 0.86 | 0.39 | 0.8565 | -0.30 | 0.78 |
| Superior Parietal Lobe L | 0.12 | 0.42 | 0.29 | 0.77 | 0.2878 | -0.70 | 0.95 |
| Superior Parietal Lobe R | 0.20 | 0.36 | 0.57 | 0.57 | 0.5723 | -0.49 | 0.90 |
| **Superior Temporal Gyrus R** | **0.41** | **0.30** | **1.37** | **0.17** | **1.3665** | **-0.18** | **0.99** |

Note: Regions with significant memorability effects are marked in bold.

### Multilevel mediation analyses

#### Encyclopedic factors and conceptual memorability

Multilevel mediation analyses showed that in several of the above ROIs showing sensitivity to object memorability, a substantial proportion of the relationship between memorability and brain activity was mediated by semantic factors. While the success of a mediation model can be defined in many different ways, we focus on regions that demonstrated both significant memorability effects in our previous analyses and significant mediation effects in the current analysis. **Table 2** and **Table 3** list the results from the mediation analyses that showed significant mediation effects, determined by 95% confidence intervals that do not include zero. Here the encyclopedic factors mediate the relationship between conceptual memorability and stimulus-specific brain activity.

**Table 3.**
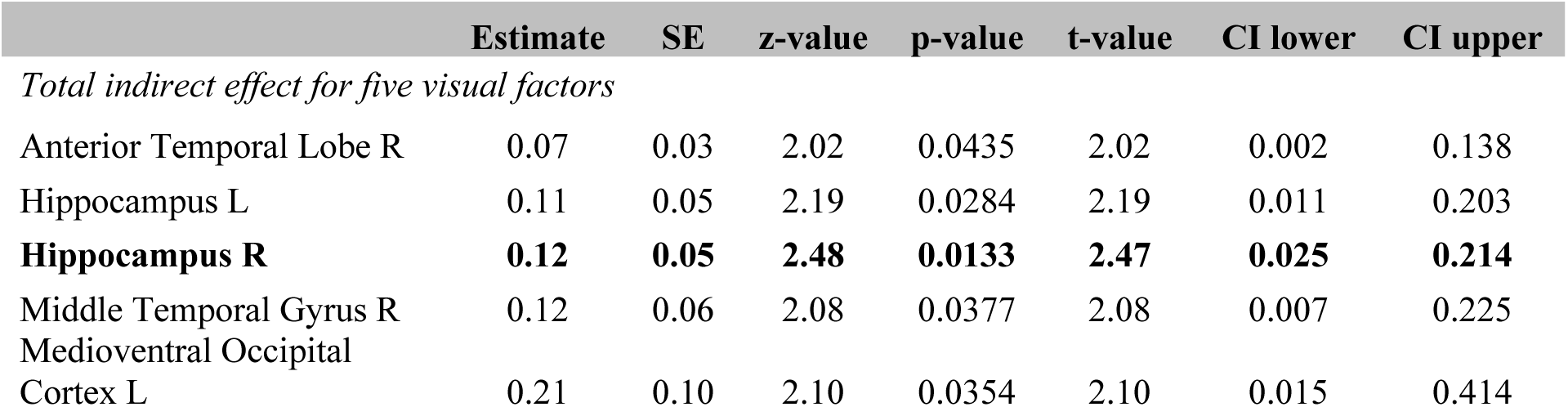
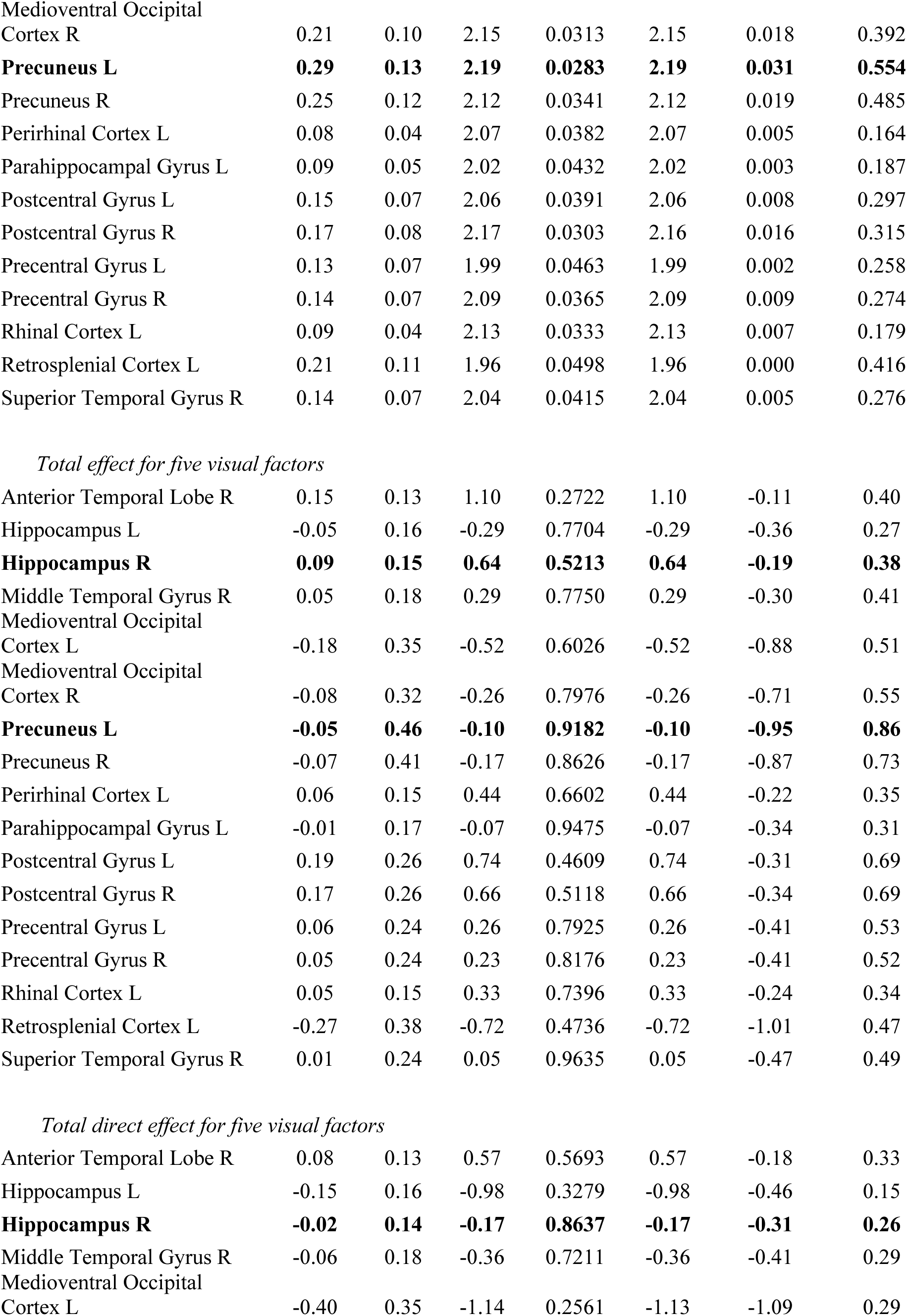
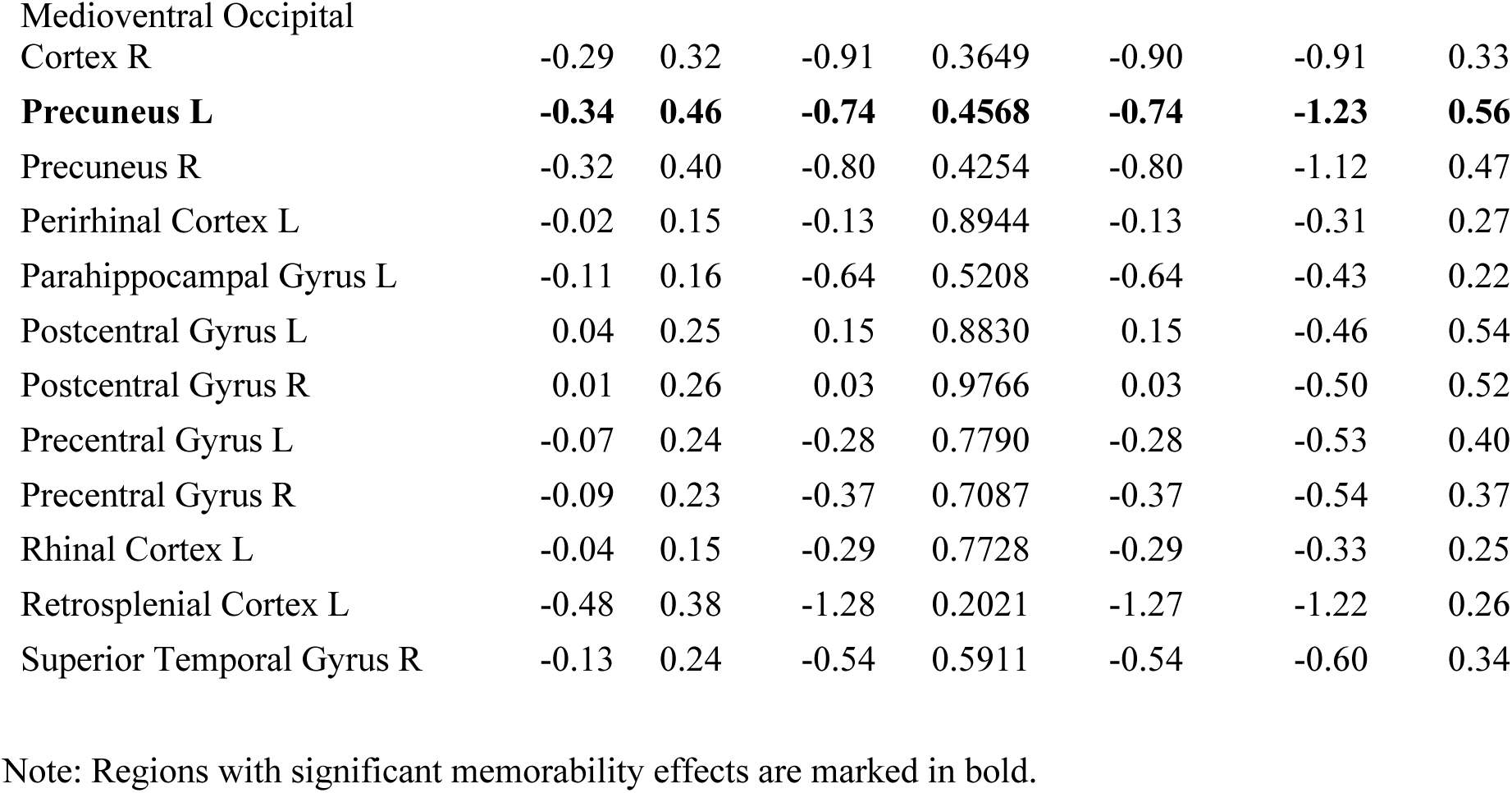
Mediation results for visual factors

|  | Estimate | SE | z-value | p-value | t-value | CI lower | CI upper |
| --- | --- | --- | --- | --- | --- | --- | --- |
| *Total indirect effect for five visual factors* |  |  |  |  |  |  |  |
| Anterior Temporal Lobe R | 0.07 | 0.03 | 2.02 | 0.0435 | 2.02 | 0.002 | 0.138 |
| Hippocampus L | 0.11 | 0.05 | 2.19 | 0.0284 | 2.19 | 0.011 | 0.203 |
| **Hippocampus R** | **0.12** | **0.05** | **2.48** | **0.0133** | **2.47** | **0.025** | **0.214** |
| Middle Temporal Gyrus R | 0.12 | 0.06 | 2.08 | 0.0377 | 2.08 | 0.007 | 0.225 |
| Medioventral Occipital Cortex L | 0.21 | 0.10 | 2.10 | 0.0354 | 2.10 | 0.015 | 0.414 |
| Medioventral Occipital Cortex R | 0.21 | 0.10 | 2.15 | 0.0313 | 2.15 | 0.018 | 0.392 |
| **Precuneus L** | **0.29** | **0.13** | **2.19** | **0.0283** | **2.19** | **0.031** | **0.554** |
| Precuneus R | 0.25 | 0.12 | 2.12 | 0.0341 | 2.12 | 0.019 | 0.485 |
| Perirhinal Cortex L | 0.08 | 0.04 | 2.07 | 0.0382 | 2.07 | 0.005 | 0.164 |
| Parahippocampal Gyrus L | 0.09 | 0.05 | 2.02 | 0.0432 | 2.02 | 0.003 | 0.187 |
| Postcentral Gyrus L | 0.15 | 0.07 | 2.06 | 0.0391 | 2.06 | 0.008 | 0.297 |
| Postcentral Gyrus R | 0.17 | 0.08 | 2.17 | 0.0303 | 2.16 | 0.016 | 0.315 |
| Precentral Gyrus L | 0.13 | 0.07 | 1.99 | 0.0463 | 1.99 | 0.002 | 0.258 |
| Precentral Gyrus R | 0.14 | 0.07 | 2.09 | 0.0365 | 2.09 | 0.009 | 0.274 |
| Rhinal Cortex L | 0.09 | 0.04 | 2.13 | 0.0333 | 2.13 | 0.007 | 0.179 |
| Retrosplenial Cortex L | 0.21 | 0.11 | 1.96 | 0.0498 | 1.96 | 0.000 | 0.416 |
| Superior Temporal Gyrus R | 0.14 | 0.07 | 2.04 | 0.0415 | 2.04 | 0.005 | 0.276 |
| *Total effect for five visual factors* |  |  |  |  |  |  |  |
| Anterior Temporal Lobe R | 0.15 | 0.13 | 1.10 | 0.2722 | 1.10 | -0.11 | 0.40 |
| Hippocampus L | -0.05 | 0.16 | -0.29 | 0.7704 | -0.29 | -0.36 | 0.27 |
| **Hippocampus R** | **0.09** | **0.15** | **0.64** | **0.5213** | **0.64** | **-0.19** | **0.38** |
| Middle Temporal Gyrus R | 0.05 | 0.18 | 0.29 | 0.7750 | 0.29 | -0.30 | 0.41 |
| Medioventral Occipital Cortex L | -0.18 | 0.35 | -0.52 | 0.6026 | -0.52 | -0.88 | 0.51 |
| Medioventral Occipital Cortex R | -0.08 | 0.32 | -0.26 | 0.7976 | -0.26 | -0.71 | 0.55 |
| **Precuneus L** | **-0.05** | **0.46** | **-0.10** | **0.9182** | **-0.10** | **-0.95** | **0.86** |
| Precuneus R | -0.07 | 0.41 | -0.17 | 0.8626 | -0.17 | -0.87 | 0.73 |
| Perirhinal Cortex L | 0.06 | 0.15 | 0.44 | 0.6602 | 0.44 | -0.22 | 0.35 |
| Parahippocampal Gyrus L | -0.01 | 0.17 | -0.07 | 0.9475 | -0.07 | -0.34 | 0.31 |
| Postcentral Gyrus L | 0.19 | 0.26 | 0.74 | 0.4609 | 0.74 | -0.31 | 0.69 |
| Postcentral Gyrus R | 0.17 | 0.26 | 0.66 | 0.5118 | 0.66 | -0.34 | 0.69 |
| Precentral Gyrus L | 0.06 | 0.24 | 0.26 | 0.7925 | 0.26 | -0.41 | 0.53 |
| Precentral Gyrus R | 0.05 | 0.24 | 0.23 | 0.8176 | 0.23 | -0.41 | 0.52 |
| Rhinal Cortex L | 0.05 | 0.15 | 0.33 | 0.7396 | 0.33 | -0.24 | 0.34 |
| Retrosplenial Cortex L | -0.27 | 0.38 | -0.72 | 0.4736 | -0.72 | -1.01 | 0.47 |
| Superior Temporal Gyrus R | 0.01 | 0.24 | 0.05 | 0.9635 | 0.05 | -0.47 | 0.49 |
| *Total direct effect for five visual factors* |  |  |  |  |  |  |  |
| Anterior Temporal Lobe R | 0.08 | 0.13 | 0.57 | 0.5693 | 0.57 | -0.18 | 0.33 |
| Hippocampus L | -0.15 | 0.16 | -0.98 | 0.3279 | -0.98 | -0.46 | 0.15 |
| **Hippocampus R** | **-0.02** | **0.14** | **-0.17** | **0.8637** | **-0.17** | **-0.31** | **0.26** |
| Middle Temporal Gyrus R | -0.06 | 0.18 | -0.36 | 0.7211 | -0.36 | -0.41 | 0.29 |
| Medioventral Occipital Cortex L | -0.40 | 0.35 | -1.14 | 0.2561 | -1.13 | -1.09 | 0.29 |
| Medioventral Occipital Cortex R | -0.29 | 0.32 | -0.91 | 0.3649 | -0.90 | -0.91 | 0.33 |
| **Precuneus L** | **-0.34** | **0.46** | **-0.74** | **0.4568** | **-0.74** | **-1.23** | **0.56** |
| Precuneus R | -0.32 | 0.40 | -0.80 | 0.4254 | -0.80 | -1.12 | 0.47 |
| Perirhinal Cortex L | -0.02 | 0.15 | -0.13 | 0.8944 | -0.13 | -0.31 | 0.27 |
| Parahippocampal Gyrus L | -0.11 | 0.16 | -0.64 | 0.5208 | -0.64 | -0.43 | 0.22 |
| Postcentral Gyrus L | 0.04 | 0.25 | 0.15 | 0.8830 | 0.15 | -0.46 | 0.54 |
| Postcentral Gyrus R | 0.01 | 0.26 | 0.03 | 0.9766 | 0.03 | -0.50 | 0.52 |
| Precentral Gyrus L | -0.07 | 0.24 | -0.28 | 0.7790 | -0.28 | -0.53 | 0.40 |
| Precentral Gyrus R | -0.09 | 0.23 | -0.37 | 0.7087 | -0.37 | -0.54 | 0.37 |
| Rhinal Cortex L | -0.04 | 0.15 | -0.29 | 0.7728 | -0.29 | -0.33 | 0.25 |
| Retrosplenial Cortex L | -0.48 | 0.38 | -1.28 | 0.2021 | -1.27 | -1.22 | 0.26 |
| Superior Temporal Gyrus R | -0.13 | 0.24 | -0.54 | 0.5911 | -0.54 | -0.60 | 0.34 |

Note: Regions with significant memorability effects are marked in bold.

Importantly, three of the ROIs that showed memorability effects also show mediation effects for encyclopedic factors. Estimates of the total indirect effect were 0.161 (*p* = 0.013) for right hippocampus, 0.411 (*p* = 0.022) for left precuneus, and 0.125 (*p* = 0.0185) for right rhinal cortex. Visual factors and factors derived from the original feature matrix mediated the relationship between perceptual memorability and activity in right hippocampus. The proportion mediated of these regions are 0.42 for right hippocampus, 0.37 for left precuneus, and 0.39 for right rhinal cortex, defined as the total indirect effect divided by the total effect (Figure 8). In the case of visual factors, right hippocampus, left precuneus, and right superior temporal gyrus showed significant total indirect effects. Functional factors did not show any significant mediation effects.

**Figure 8.**
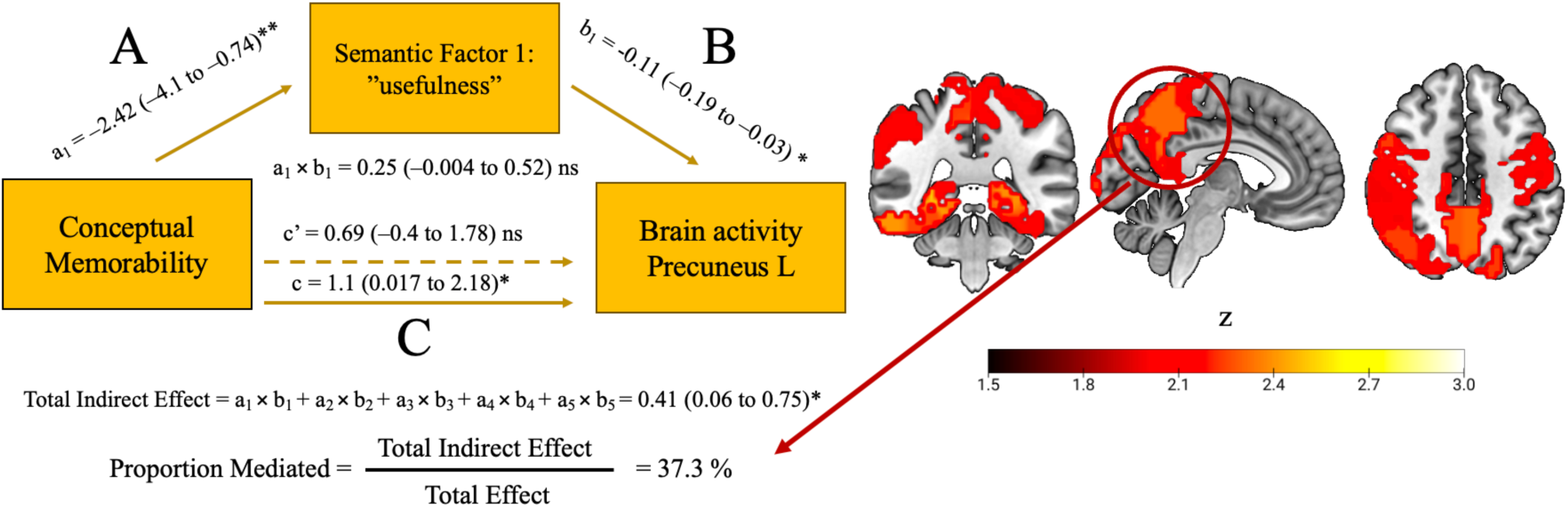
Sample mediation for ROI with highest proportion mediated. Complete mediation analysis and calculation of proportion mediated for one region. In this case, the relationship between stimulus memorability and activity in left precuneus is mediated by the first encyclopedic factor, characterized by the top feature “is useful.” The proportion mediated is the quotient of the total indirect effect and total effect.

Interestingly, while 28 regions showed significant total indirect effects for encyclopedic factors, 13 of these had no significant individual indirect effects, meaning that the indirect effects of individual factors alone were not significant. On the other hand, 15 regions showed a significant indirect effect of Factor 1, and no other factor was individually significant (**Table 4**). This effect may be explained by the fact that NMF generates factors which capture semantics in an aggregate manner. While individual factors may not be significant mediators on their own, they operate as significant mediators of the relationship between stimulus memorability and stimulus-specific brain activity as a group.

**Table 4.**
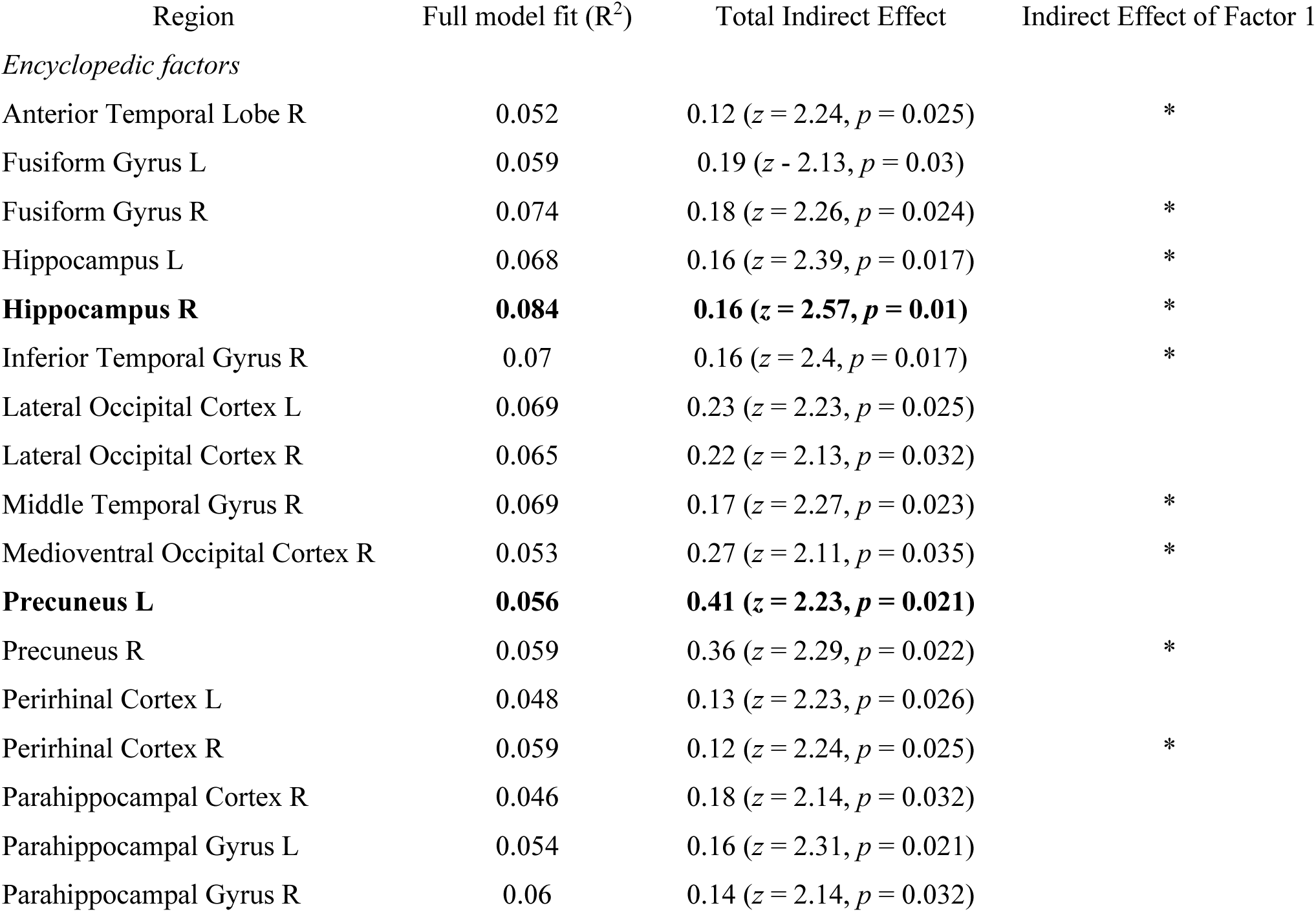
Individual vs total indirect effects

**Table 4.**
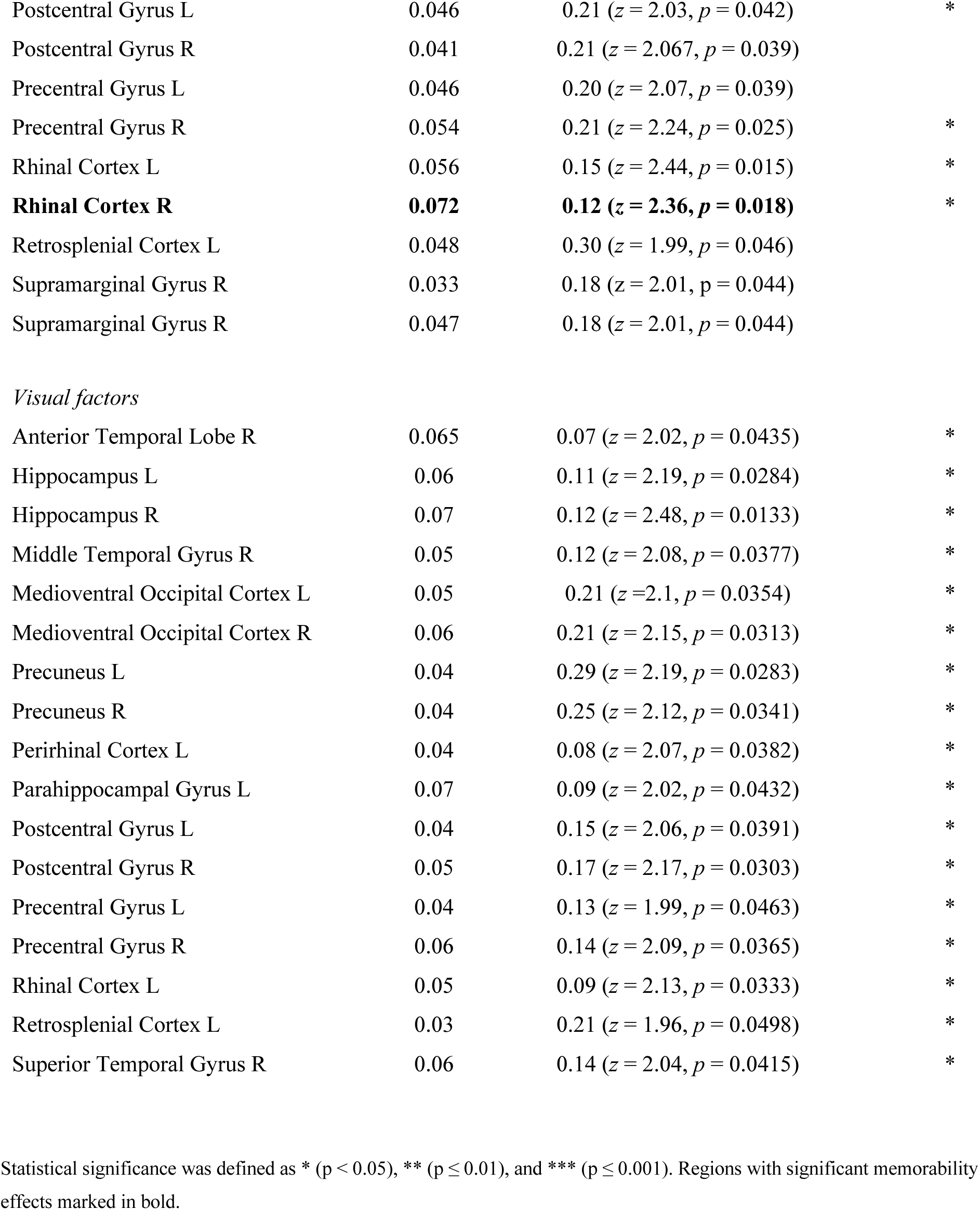
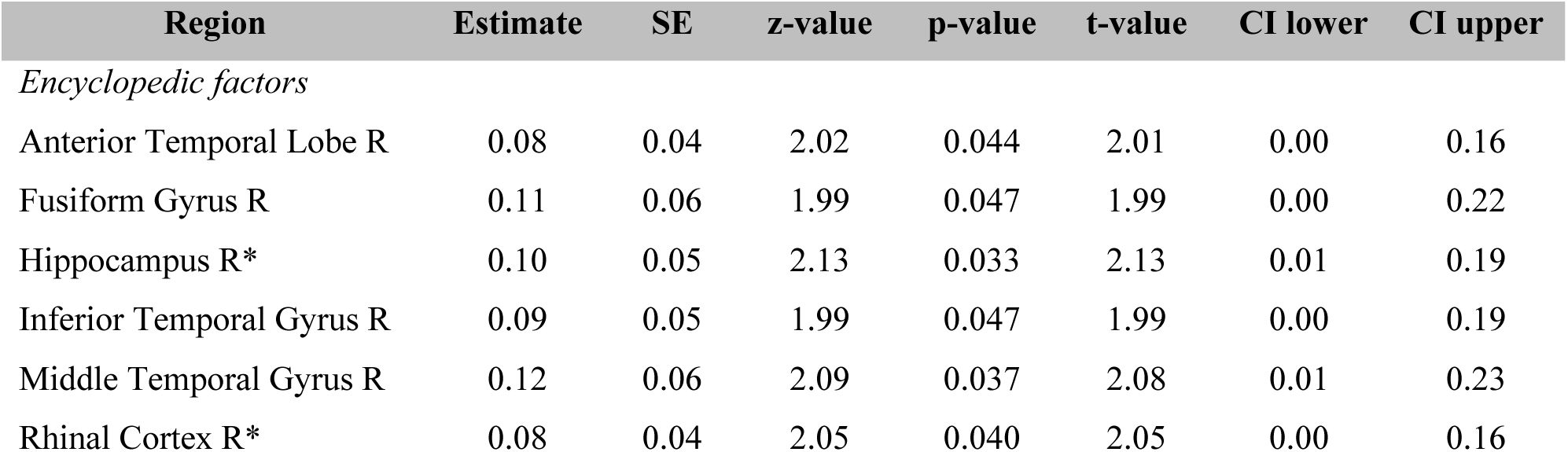
Single-mediator results

This effect can be seen by comparing the multiple-mediator model with a single-mediator model. We ran an additional mediation model using only encyclopedic Factor 1 and there were fewer significant regions (**Table 4**). Interestingly, while right hippocampus and right rhinal cortex showed a significant indirect effect, left precuneus did not. This would suggest that semantic factors only mediate the memorability effect in that region as a group. Multiple mediation has several advantages over single-mediator models, not only for assessing the presence of an overall effect, but for comparing the relative importance of each mediator and reducing omitted variable bias (Preacher & Hayes, 2008). It may be that the total indirect effect is significant and that individual indirect effects are not, or the total effect is not significant and the individual effects are, which is why it is important to consider the differences between the model approaches.

In this case, we can infer that Factor 1 (i.e., “usefulness”) is especially important given that it is the only factor that is a significant mediator on its own. In the case of visual factors, all regions with significant total indirect effects also showed a significant indirect effect for Factor 1, but not for other individual factors. The feature, “made of metal” is an important differentiating factor for the visual appearance of the stimuli. It may be that utility or material are so important for the mediating effect of semantic factors because they are general semantic properties that apply to our entire data set of concrete objects. They are characteristic of the objects whether they are useful or not or made of metal or not, and therefore may serve a central conceptual role in how they are perceived and remembered. When considering the feature norms from which the factors are derived, we can see that “is useful” (the top-loading feature on encyclopedic Factor 1) is correlated with several other features, including “is helpful,” “is practical,” and “is needed.” Similarly, the top-loading feature on visual Factor 1 “is made of metal” is correlated with other material-based features, such as “is made of steel,” “is shiny,” and “is silver” (Figure 6). While factors with a high number of mentions in the feature norm data set (such as “is useful” and “is made of metal”) load most highly on each factor, we can see that they are representative of conceptual clusters within the feature matrix. Because the factors characterized by utility and material are significant mediators on their own, we can infer that they are important for the memorability effect on stimulus-specific brain activity during encoding. We can further infer that because for many ROIs the five factors are significant mediators as a group though not individually, that they capture complementary aspects of the semantics of the stimuli, and that these object feature semantics mediate a substantial portion of the relationship between memorability and brain activity.

The strongest mediation effects were observed in semantic (e.g. Precuneus, Inferior Parietal Lobule), motor (Precentral Gyrus, Postcentral Gyrus), and visual regions (Medioventral occipital cortex, lateral occipital cortex) (Figure 9). Several studies have observed that the representation of objects involves a kind of embodiment, where perception, action, and mode-specific activity support the encoding and retrieval of those objects (Binder et al., 2016; Fernandino et al., 2015, 2022; Tong et al., 2022). In this way, objects are differentially memorable when brain activity in particular regions is stronger. Importantly, we see a positive relationship, suggesting that increased activity in these regions is associated with enhanced object memorability.

**Figure 9.**
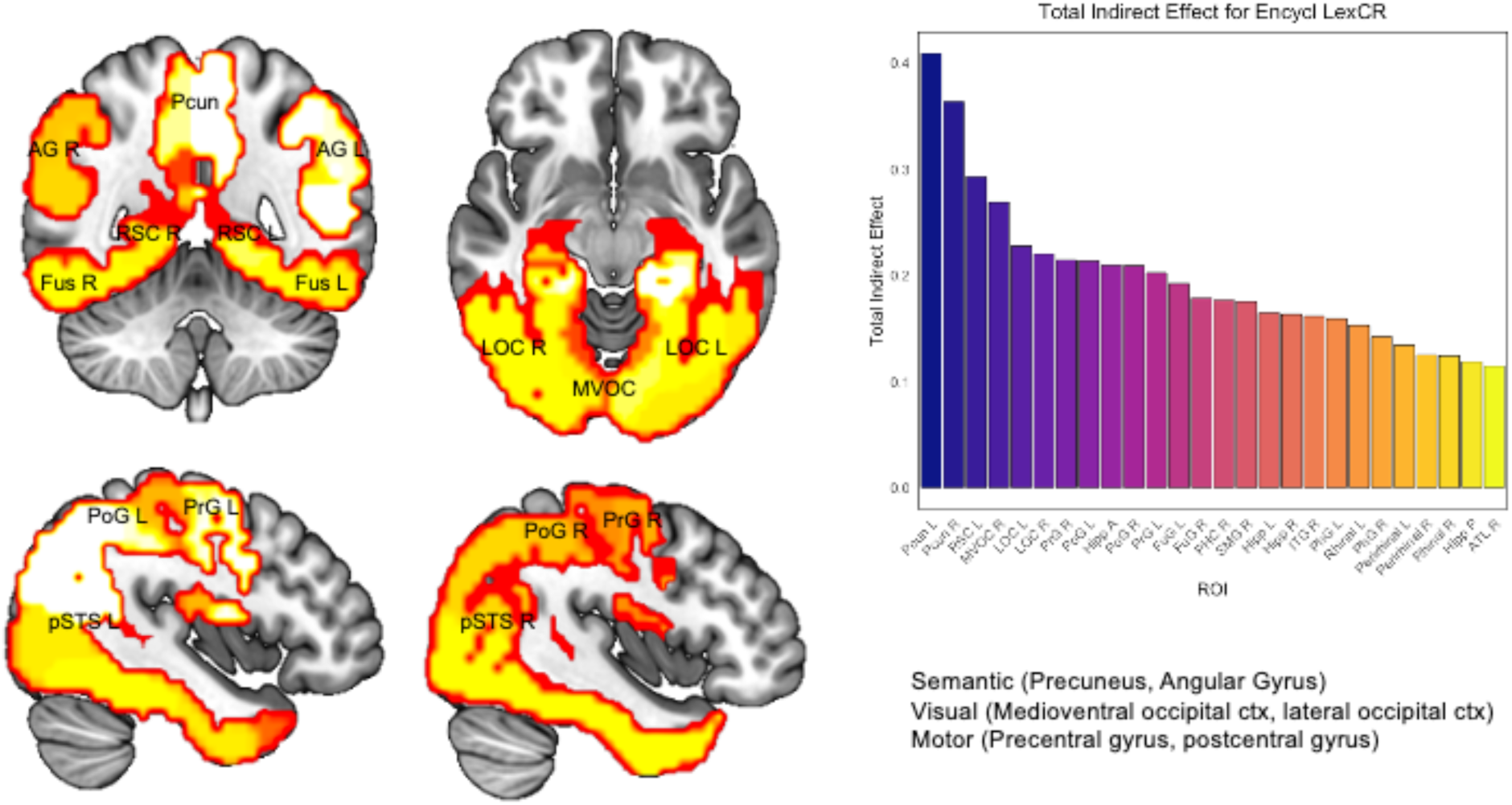
Sensory, motor, and visual regions show the strongest mediation effects. Representative figure showing the total indirect effects from mediation models exploring the effect of the top five semantic factors on the relationship between memorability and activity in each region. The regions with the highest total indirect effects fall into three categories, supporting semantic, visual, and motor processes.

## DISCUSSION

The current study sought to use a broad empirical approach to evaluate determinant factors for the neural basis of object memorability. First, we identified an object-wise effect of memorability in rhinal cortex and hippocampus, ventral temporal regions known to play a role in the memory and conceptual processing. Second, we found that semantic factors play an important role in the relationship between memorability and brain activity, mediating up to 42% of the relationship between memorability and brain activity in right hippocampus, 39% in right rhinal cortex, and 37% in left precuneus (as defined by the total indirect effect divided by the total effect). Importantly, the regions that show the strongest mediation effect perform semantic, visual, or motor functions. These results offer a more complete view on how semantic information mediates the relationship between memorability and brain activity.

Our first principal finding is that multiple mnemonic regions in the temporal lobe (hippocampus, rhinal cortex) demonstrated significant relationships between BOLD activation and object-wise memorability. In addition to these canonical regions important for episodic memory encoding (Danieli et al., 2023), we found significant associations with memorability in the superior temporal gyrus, a region involved in language and semantic processes that support memory (Liu et al., 2023), as well as the left precuneus and right inferior parietal lobule, regions associated with semantic memory processing (Binder & Desai, 2011). Activity in precuneus and inferior parietal lobule in particular were associated with predicted conceptual memorability, consistent with the role these regions play in the integration of episodic and semantic information (Murphy et al., 2023).

The concept of memorability as an *intrinsic attribute of images* arose in several studies that sought to identify factors that affect memory. For example, studies of memory for faces found that distinctiveness and typicality (Valentine, 1991) and prototypicality (Cabeza et al., 1999) can affect how memorable a given face is. And yet, later studies suggest that memorability may be an intrinsic property because individual traits like typicality or trustworthiness cannot adequately account for the phenomenon (Bainbridge et al., 2013). What’s more, memorability estimates are generally quite reliable across random populations (Isola et al., 2014), though these estimates are sensitive to particularities of the dataset and the individual images (Khosla et al., 2015). The memorability effects, though modest, help connect trial-level variation in mean brain activity with more traditional analyses that rely on condition-level variation to understand subsequent memory in the hippocampus and rhinal cortex (Fernández et al., 1999).

Our second principal finding is that semantic factors, derived from nonnegative matrix factorization applied to image features, appear to play an important role in mediating the relationship between memorability and brain activity. Furthermore, individual dimensions moderate the relationship between memorability and brain activity (**Supplementary Table 1**), whereas the magnitude of the loading of an image on a particular dimension determines whether memorability drives brain activity in a particular region. In the case of encyclopedic factors, we saw that a factor characterized by usefulness was the only factor that had a significant indirect effect as an individual. For visual factors, a factor characterized by the property “made of metal” was the only individual factor with a significant indirect effect. As a group, encyclopedic factors characterized by usefulness, animacy, engagement, weapon, and marine animacy together showed a significant indirect effect in several regions, including those involved in episodic memory, semantic memory, and language.

Taken together, these results provide strong evidence for the role of semantic information in facilitating the encoding of memories in the brain. It has been long appreciated that the deeper information is processed, the longer a memory trace will last (Craik & Lockhart, 1972). Craik and Lockhart defined *depth* as the meaningfulness extracted from the stimulus, where deeper levels of processing are organized along structural, phonetic, and semantic levels. More recently, several studies have tried to quantify these semantic dimensions in order to understand their representation in the brain while participants perform perception (Contier et al., 2023; Kramer et al., 2022; Muttenthaler & Hebart, 2022) and memory tasks (Davis et al., 2021; Huang et al., 2024; Kramer et al., 2023). We argue that *memorability* can be seen as a conceptual short-hand for sensory-functional processes that facilitate the visual encoding of semantic information. Such results follow similar conclusions made by groups who have identified the representation of conceptual information in sensory-motor and affective areas (Binder et al., 2016; Fernandino et al., 2015, 2022; Tong et al., 2022), which suggests that these effects are driven by sensory and functional processes (Mahon & Caramazza, 2009). We can see evidence for such an account in the mediation analyses, where many regions showed a significant mediation effect despite not being typically associated with memory. These include regions involved in semantic, visual, and motor functions. We can infer that sensory and motor processes support the encoding and retrieval of the images. This sensory-functional account for concepts in the brain has important consequences for the question of memorability. It may be that the image-inherent property of memorability can be, in principle, decomposed into the conceptual, perceptual, and motor components of the interpretation and understanding of the objects in the images.

### Potential limitations

There are several limitations of this study, such as it being an exploratory, post hoc analysis. Furthermore, there are many ways to define and quantify semantic features. Two such methods include the feature norm approach used in this study (Hovhannisyan et al., 2021) and extracting features from similarity judgments (Hebart et al., 2020; Kramer et al., 2023). We selected feature norms given its history of use in psychology and cognitive neuroscience, but future studies might employ additional methodologies, such as corpus or database modeling (Lindh-Knuutila & Honkela, 2015), relationship extraction methods (Perez-Arriaga et al., 2018), or word-and sentence-embedding methods such as word2vec (Mikolov, Chen, et al., 2013; Mikolov, Sutskever, et al., 2013), GloVe (Pennington et al., 2014), or BERT (Devlin et al., 2018). Several authors combine approaches, such as Bhatia and Richie who use feature norms to refine a pre-trained BERT model to increase model performance (Bhatia & Richie, 2024) or feature2vec to augment participant-derived feature norm datasets (Derby et al., 2019). A systematic comparison of the similarity in semantic feature spaces derived from these disparate techniques could be especially useful for exploring the extent of the relationship between semantics and memory given the particular benefits and costs of each modeling approach.

While the object-wise approach has been employed previously (Hovhannisyan 2021; Kramer et al. 2022), task-specific conditions may affect the reliability of object-level memorability effects. Activity estimates in this study were derived from a simple object encoding task, during which participants were presented with a single letter and color image depicting an everyday object, and only pressed a button when they noticed a discrepancy between the letter and name of the presented object. **Table 1** outlines effects limited to subsequently remembered trials, a step meant to exclude trials during which participants may have not been focused on the stimuli (or there was some other interference that led to subsequent forgetting of that stimulus). Such an approach is useful for modeling subsequent memory at an object-wise level, but it may be useful to explore subsequent forgetting effects as well. Finally, future studies could employ a multivariate method such as Representational Similarity Analysis to explore how representational strength affects memorability, though similar methods have been used to find that both visual and semantic information in hippocampus and rhinal cortex contributes to subsequent conceptual and perceptual memory (Davis et al., 2021) and that representational changes depend on memory phase (i.e., encoding or retrieval) (Howard et al., 2024).

## Conclusion

In conclusion, our approach to modeling image semantics by applying NMF to feature norms identified several regions which support normative image memorability derived from a separate population. These factors mediate a large proportion of the relationship between memorability and visually-presented, image-specific brain activity. Such effects can be seen in regions involved not only in episodic memory and semantics, but in motor regions as well. Together, these findings suggest that image semantic properties can partially explain the phenomenon of memorability. It may be that image memorability can be decomposed into conceptual, perceptual, and motor components, which are in turn reflected in the processing of these images during memory encoding.

The authors declare no conflicts of interest.

## Acknowledgement

This work was supported by the National Institute of Health, R01-AG066901, R01-AG075417.

