## Supplement for "Semantic Dimensions Support the Cortical Representation of Object Memorability"

### SUPPLEMENTARY MATERIALS

#### Supplementary Methods

*Methodological considerations in applying non-negative matrix factorization*

**Thresholding**

In order to determine the appropriate number of features, we used a two-step process. First, a thresholding procedure was applied to each feature matrix, to remove feature columns below a cutoff. Using the skewness function in MATLAB, the moment coefficient of skewness was calculated for each column in each feature matrix which was in turn averaged to give a mean skewness of each matrix: 21.44 for the entire matrix, and for the subset matrices, 21.41 for encyclopedic, 20.59 for visual, 23.83 for functional. Based on the mean skewness, the baseline threshold of 50$^{th}$ percentile was dynamically adjusted to determine the threshold below which feature columns were removed in order to ensure that the most relevant features were retained. The 70.45$^{th}$ percentile was calculated for the entire matrix, yielding a threshold of features with fewer than 10 mentions in the feature norm data set. For encyclopedic, 70.41th percentile, yielding a threshold of 9, for visual 69.59$^{th}$ percentile and a threshold of 12, and for functional 72.83th percentile and a threshold of 8.

**Determining Optimal Rank**

In order to determine the optimal rank of NMF for our data, we used two methods. First, Knee Point Detection, a method by which NMF is applied with iteratively more factors, the original matrix is reconstructed from the component W and H matrices, and the error between the original and reconstructed matrix is determined. The addition of factors reduces the error between the two matrices, and the knee point is defined as the number of factors where the decrease in residuals falls below a threshold.

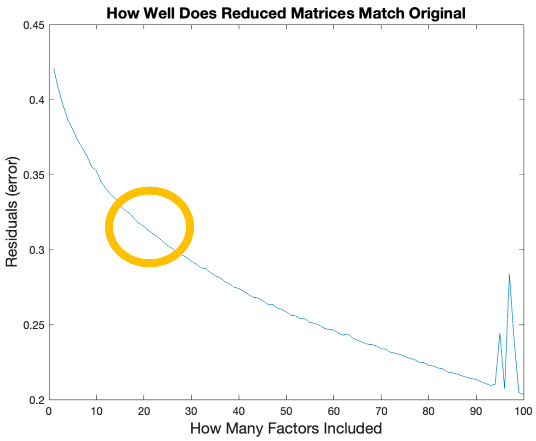


***Supplementary Figure 1. Knee Point Detection.*** One common approach to determining the optimal rank of NMF is to investigate the residual error between original and reconstruction matrix by serially adding additional factors. The knee point is the point at which the gains in model fit become smaller for each new factor added.

A second method was applied that used cross-validation to evaluate model performance and prevent over-fitting. First, imputation was used to randomly mask elements of the data in order to split the original matrix into train and test groups. NMF was fit on the training set with an iteratively increasing number of factors, and the reconstruction error was calculated based on the test data. The minimum reconstruction error determines the optimal rank. Given the agreement between these two approaches, we considered the determined rank to be valid.

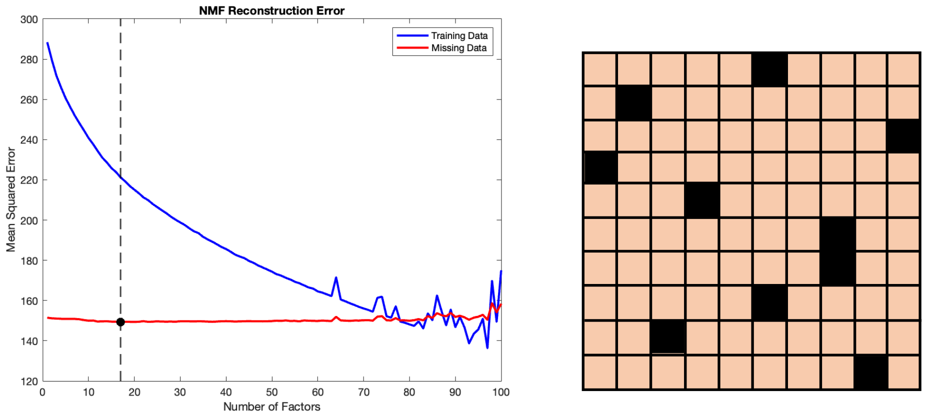


***Supplementary Figure 2.*** Another strategy for determining NMF rank is to investigate the performance of NMF on a training set with the reconstruction error based on test data. The data is split into training and testing sets by removing individual elements from the matrix (i.e. cros-validation) in order to ensure robustness and avoid bias.

**Factor Selection**

With the number of factors determined, we performed additional factor-selection procedures to determine the relative importance of each factor. Frobenius norm reconstruction error, similar to its use in the optimal rank procedure, can show the relative importance of each factor, as all but one factor is used to reconstruct the original matrix, and the higher the error between the original and reconstructed matrices, the more important the left-out factor is. We additionally applied Elastic Net regularization (a technique which combines the L1 penalty of LASSO and the L2 penalty of Ridge regression) to determine factor importance by performing variable selection and shrinking the coefficients of less important factors toward zero. Finally, we applied a procedure which involved modeling all factors as predictors and iteratively modeling all factors with one excluded. The $R^2$ for the "all factors" model was subtracted from the $R^2$ for the "one factor excluded" model to estimate the importance of the excluded factor. These factors were then sorted from most to least important in terms of "$R^2$ difference." These factors were added one at a time to a model memorability ~ factors and the adjusted $R^2$ was calculated. Then a procedure was used to determine where the 'knee' was in the cumulative adjusted $R^2$. Using a sliding window of 3, we asked where the differences fell below a threshold of .0005. This is where additional factors led to diminishing returns in terms of model fit. Betas were then calculated for each factor and significance determined with a threshold of .05.

**Supplementary Table 1**

| Region | Factor | Beta | SE | t-value | p-value | p-value adjusted | F |
|---|---|---|---|---|---|---|---|
| Inferior Frontal Gyrus L | Encycl F3 | -0.63 | 0.24 | -2.60 | 0.0100 | 0.0299 | 1.23 |
| Parahippocampal Cortex L | Encycl F5 | -1.27 | 0.54 | -2.35 | 0.0198 | 0.0394 | 3.44 |
| Parahippocampal Gyrus L | Encycl F5 | -0.97 | 0.39 | -2.51 | 0.0129 | 0.0387 | 3.95 |
| Rhinal Cortex L | Encycl F5 | -0.85 | 0.35 | -2.41 | 0.0166 | 0.0498 | 3.71 |
| Rhinal Cortex R | Encycl F1 | 0.17 | 0.07 | 2.44 | 0.0156 | 0.0467 | 0.66 |
